# Delineation and birth of a layered intestinal stem cell niche

**DOI:** 10.1101/2021.09.28.462142

**Authors:** Neil McCarthy, Guodong Tie, Shariq Madha, Judith Kraiczy, Adrianna Maglieri, Ramesh A. Shivdasani

## Abstract

Wnt and Rspondin (RSPO) signaling drives proliferation, and bone morphogenetic protein inhibitors (BMPi) impede differentiation, of intestinal stem cells (ISCs). Here we identify the adult mouse ISC niche as a complex, multi-layered structure that encompasses distinct mesenchymal and smooth muscle cell populations. Diverse sub-cryptal cells provide redundant supportive factors, with certain BMPi and the most potent Wnt co-factor, RSPO2, restricted to single cell types. Niche functions refine during a critical period of postnatal crypt morphogenesis, in part to oppose dense aggregation of BMP-expressing sub-epithelial myofibroblasts that promote epithelial differentiation. A specialized muscle layer, the muscularis mucosae, first appears during this period and supplements neighboring RSPO and BMPi sources. In vivo ablation of smooth muscle raises BMP activity and potently limits a pre-weaning burst of crypt fission. Thus, distinct and progressively specialized mesenchymal components together create the milieu required to propagate crypts during rapid organ growth and to sustain adult ISCs.

## Introduction

Multipotent *Lgr5^+^* intestinal stem cells (ISCs) proliferate at the base of adult small intestine (SI) crypts (Barker et al., 2007), relinquishing stem-cell properties as they leave that niche (Ritsma et al., 2014). Epithelial homeostasis balances Wnt/RSPO-dependent ISC self-renewal (Clevers, 2013) against bone morphogenetic protein (BMP)-mediated differentiation (Qi et al., 2017). Hence, mutations in human BMP signaling genes *SMAD4*, *BMPR1A* or *GREM1* expand ISCs (Howe et al., 2001; Howe et al., 1998; Jaeger et al., 2012) and forced BMPi expression in mice induces ectopic crypts (Batts et al., 2006; Davis et al., 2015; Haramis et al., 2004). The crypt base is a zone of high Wnt/RSPO and low BMP activity, the same milieu required to expand ISCs *ex vivo* (Sato et al., 2009). Sub-epithelial tissue is the principal source of these signals (Farin et al., 2012; McCarthy et al., 2020a).

Mouse villus morphogenesis has been elegantly described (Walton et al., 2016) and fetal colon cell types were recently catalogued (Brugger et al., 2020). At birth, however, the mouse SI lacks crypts. Over the ensuing ∼2 weeks (Calvert and Pothier, 1990; Itzkovitz et al., 2012), crypts progressively lengthen (Sumigray et al., 2018a), sequester ISCs at the bottom alongside Paneth cells, and later undergo fission as a dominant mode of epithelial expansion (Cheng and Bjerknes, 1985; Langlands et al., 2016; Maskens and Dujardin-Loits, 1981; St Clair and Osborne, 1985). Crypt size (Al-Nafussi and Wright, 1982), bifurcation (Cheng and Bjerknes, 1985; St Clair and Osborne, 1985), monoclonality (Schmidt et al., 1988), and Paneth cell numbers (Bry et al., 1994) increase dramatically in the third week of life. The period of substantial epithelial and ISC expansion between birth and weaning (3 weeks) is likely associated with adjacent mesenchymal sculpting.

This mesenchyme, historically regarded as a collection of myofibroblasts (Powell et al., 2011), has come into sharp focus, first through identification of overlapping CD34^+^Gp38^+^ and PDGFRA^+^ cell sources of crucial niche factors (Greicius et al., 2018; Kim et al., 2020; Kosinski et al., 2007; Stzepourginski et al., 2017). PDGFRA^hi^ cells abutting the epithelium resemble telocytes, diverse cells with a distinctive morphology found in many organs (Cretoiu et al., 2012; Popescu and Faussone-Pellegrini, 2010), and are proposed as a key source of canonical Wnt ligands (Shoshkes-Carmel et al., 2018). These cells are the same as intestinal sub-epithelial myofibroblasts (ISEMFs) long described in the literature (Powell et al., 2005; Roulis and Flavell, 2016), and because telocytes lack a defined molecular or functional identity across tissues, here we use the traditional ISEMF nomenclature. Sub-cryptal PDGFRA^lo^ cells that co-express the BMPi *Grem1* and *Rspo3* support ISC expansion *ex vivo* in the absence of exogenous trophic factors and are therefore called trophocytes (McCarthy et al., 2020b). Smooth muscle (SM) and other PDGFRA^lo^ cells are found in the trophocyte vicinity, but the totality of mesenchymal cells that serve niche functions or how the niche arises remain unclear; sub-cryptal SM, for example, also expresses *Grem1* (Koppens et al., 2021; Martin-Alonso et al., 2021).

Here we identify adult SM contributions toward the ISC niche and show that distinct SM populations substitute in part for selected trophic factors in *ex vivo* ISC cultures. The niche matures in parallel with postnatal murine crypts, coincident with increasing ISEMF density at the villus base. Niche elements positioned to counteract the resulting concentrated BMP source include the muscularis mucosae (MM) and previously uncharacterized superficial circular SM cells in the muscularis propria (MP). MM is molecularly distinct from physically contiguous lamina propria SM, expresses high *Rspo3* and *Grem2*, and arises *de novo* from resident stromal cells. It delimits a distinct anatomic compartment for trophocytes, which alone provide the most potent RSPO ligand, RSPO2. SM ablation *in vivo* during the pre-weaning peak in crypt fission augments local BMP activity and markedly attenuates crypt bifurcation. Thus, a partially redundant multi-layer ISC niche develops step-wise, as distinct mesenchymal cell types express overlapping trophic factors at defined distances from ISCs at the crypt base.

## Results

### Formation and activity of the isthmus ISEMF shelf, a potent BMP source

BMP signaling, which induces intestinal epithelial maturation (Chen et al., 2019; Qi et al., 2017), is restricted to adult mouse villi; accordingly, phospho-SMAD1/5 (pSMAD) is absent in crypt cells (Haramis et al., 2004). Epithelial pSMAD staining at postnatal days (P) 1 and 5 is confined to the top halves of emerging and enlarging villi, becoming steadily more pronounced at villus bottoms by P14 (Figures 1A and S1A, note: pSMAD signal is generally stronger in the stroma). This base-directed progression of epithelial BMP signaling could reflect increased ligand exposure or increased epithelial responsiveness; we therefore tracked the appearance of ISEMFs, which are the dominant adult source of BMP transcripts and concentrate at villus tips and bases (Bahar Halpern et al., 2020; Brugger et al., 2020; McCarthy et al., 2020b). In *Pdgfra^H2B-eGFP^* mice (Hamilton et al., 2003) (see Table S1), PDGFRA^hi^ fetal ISEMF precursors constitute the leading edges of new villi (Karlsson et al., 2000; Walton et al., 2016) (Figure S1B). These GFP^hi^ cells underlie the villus epithelium after birth, and through at least the first week of life, high ISEMF density is evident only at villus tops (Figure 1B). Aggregation of ISEMFs at the villus base was negligible at P1 and, quantified relative to the villus trunk, progressively increased through P21 (Figure 1C). Thus, the villus epithelial BMP response reflected in nuclear pSMAD staining strengthens in parallel with ISEMF density over a prominent shelf at the crypt-villus junction (isthmus – Figure 1D).

**Figure 1.**
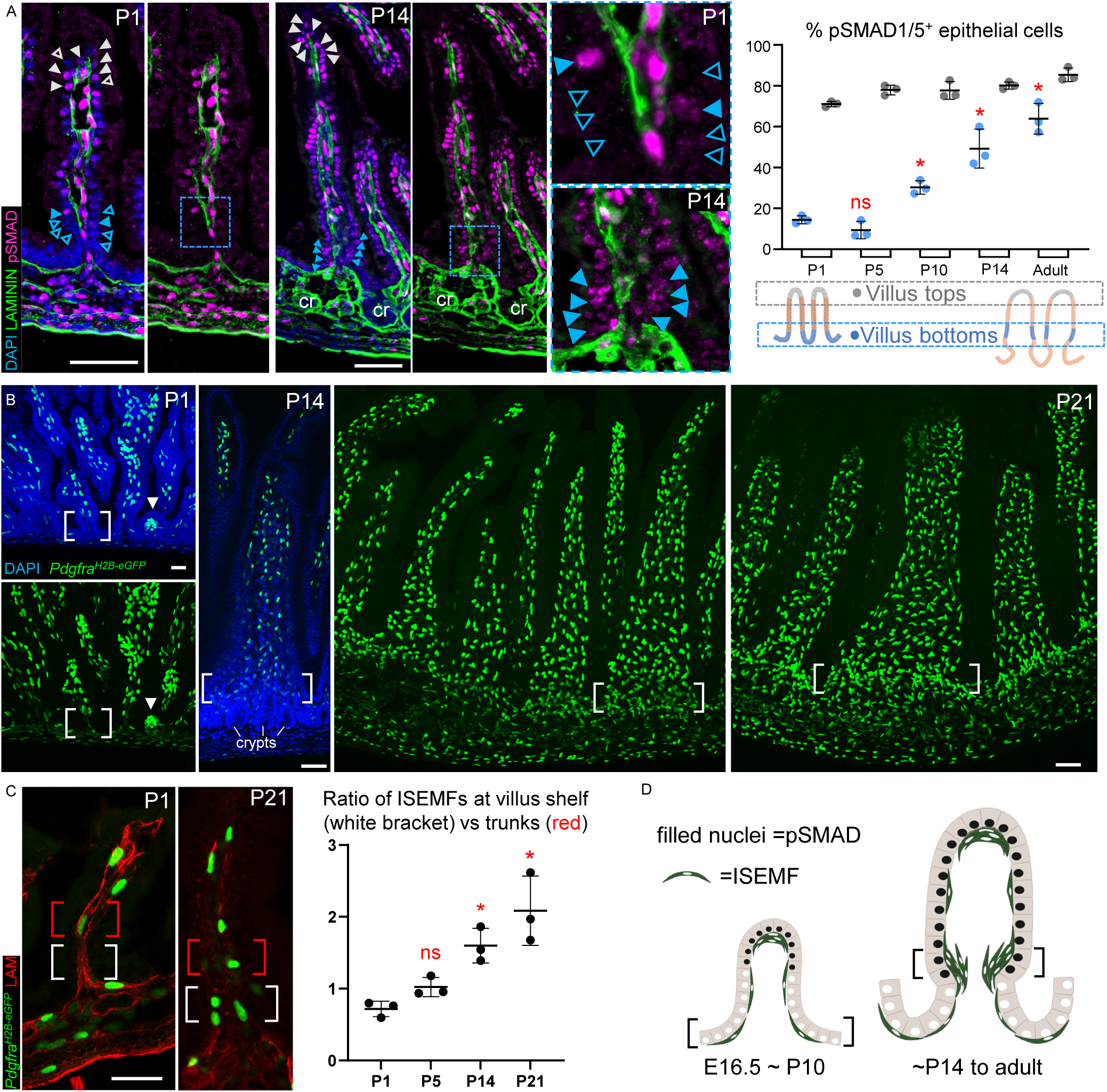
Postnatal epithelial BMP signaling coincides with isthmus ISEMF aggregation. **A)** Representative pSMAD1/5 (magenta) immunostaining (green: Laminin/basement membrane, blue: DAPI) in mouse proximal SI on indicated postnatal days. cr, crypts; scale bars 50 μm; boxed regions are magnified in adjoining images; extended images are shown in Figure S1A. Nuclear pSMAD^+^ epithelial cells (filled arrowheads; pSMAD^-^ cells indicated by empty arrowheads) are present at villus tops from birth, while pSMAD^+^ cells at villus bottoms increase over the ensuing 2 weeks. Graph displays pSMAD1/5^+^ epithelial cell fractions at the bottoms (blue dots) and tops (grey dots) of >25 villi/sample (n = 3 mice at each age). Statistical differences are reported relative to P1 villus bottoms after two-way ANOVA followed by Tukey’s multiple comparisons test. ns: not significant, *p<0.0001. **B-C)**, GFP^hi^ ISEMFs are present in *Pdgfra^H2B-eGFP^* mice on established and emerging (arrowhead) villi at P1 and concentrate progressively at the crypt-villus isthmus shelf (white brackets). From representative images of proximal SI at the indicated ages (**B**, scale bars 50 μm) we quantified GFP^hi^ ISEMF ratios (**C**) at the isthmus (white brackets, 5 lowest villus epithelial cells) vs. villus trunks (red brackets, next 5 cells toward villus tip); graph displays data from >25 villi/sample (n=3 animals at each age). Statistical differences are reported relative to P1 after two-way ANOVA followed by Tukey’s multiple comparisons test. ns: not significant, *p<0.0001. **D)** Increased epithelial pSMAD1/5 is concomitant with increasing isthmus ISEMF density. See also Figure S1.

To test whether adult ISEMFs are a functional BMP source, as mRNA profiles indicate (Brugger et al., 2020; McCarthy et al., 2020b), we cultured adult mouse duodenal crypts in media containing recombinant (r) epidermal growth factor (EGF), rRSPO1, and the BMPi rNOG (together abbreviated ENR). After organoids had established, we replaced rNOG with various factors or with stromal cell types isolated by flow cytometry (Figures 2A and S1C). Without exogenous cells or in the presence of GFP^-^ cells from *Pdgfra^H2B-eGFP^* mice (PDGFRA^-^ cells), organoids enlarged into budding structures that carry ISCs (Sato et al., 2009); in contrast, exposure to rBMPs or to GFP^hi^ ISEMFs from adult (>8 weeks old) or P14 mice arrested organoid growth. Surviving organoids retracted the classic buds (Figure S1D), which carry ISCs (Sato et al., 2009) and take up the S-phase marker 5-ethynyl-2’-deoxyuridine (EdU), and became unbranched or spheroidal; exposure to BMPi rNOG or rGREM1 restored budding morphology (Figures 2A-B and S1E). Subtle morphologic differences between organoids exposed to rBMPs or ISEMFs may reflect disparate BMP levels or other ISEMF factors, e.g., non-canonical Wnts. Organoids co-cultured with ISEMFs reduced expression of ISC molecular markers and elevated expression of BMP target genes and differentiation markers (Figure 2C). Thus, ISEMFs tilt the balance from ISC self-renewal to differentiation, in line with their post-natal aggregation at the villus base.

**Figure 2.**
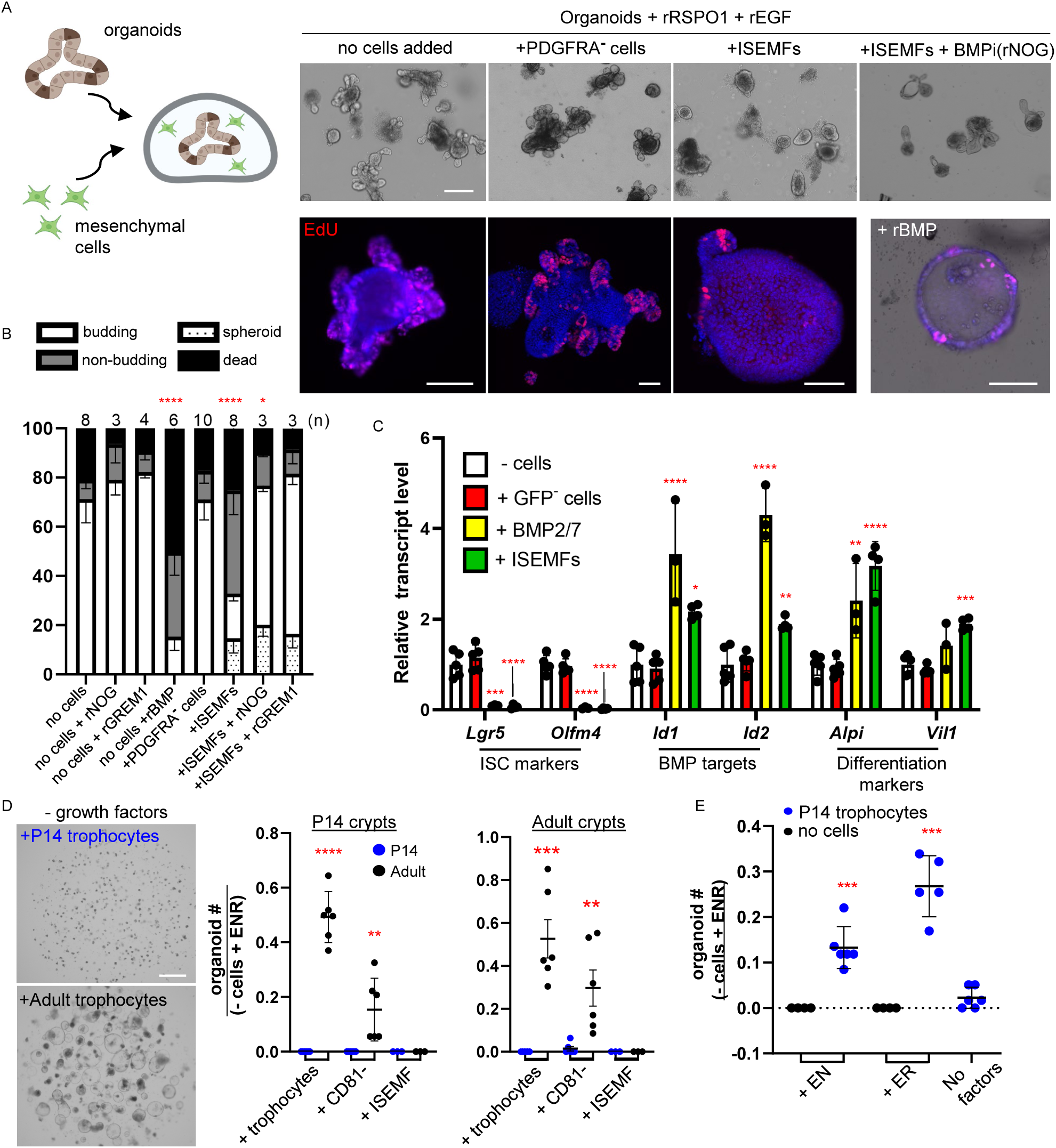
ISEMFs promote differentiation and early postnatal trophocytes fail to support organoid growth. **A)** 2 day-old organoids cultured with recombinant (r) RSPO1, EGF, and indicated cell types from *Pdgfra^H2B-eGFP^* mouse SI. Representative organoids are shown 48 h later (scale bar 100 μm). Co-cultures without cells or with GFP^-^(i.e. PDGFRA-) cells produced budding structures, whereas crypts co-cultured with ISEMFs or rBMPs retracted buds, a process that BMP inhibitor rNOG or rGREM1 reversed. Representative images of EdU-stained organoids (scale bar 50 μm) demonstrate markedly reduced cell replication in organoids co-cultured with ISEMFs or rBMPs. **B)** Fractions of budding (white), unbranched (non-budding, grey), fully spheroidal (stippled), and non-viable (black) organoids after co-culture with the indicated cells or factors. Statistics were determined for budding fractions by one-way ANOVA followed by Dunnett’s posttest. *p<0.05, ****p<0.0001 (n=3-10 independent organoid cultures as indicated above each bar). **C)** qRT-PCR analysis of organoids after 48 h co-culture (n=3 or 4 biological replicates). BMP- or ISEMF-exposed organoids reduced expression of ISC markers and increased markers of BMP activation and epithelial differentiation. Transcript levels are represented relative to control crypt cultures with no added cells. Statistical comparisons use one-way ANOVA followed by Dunnett’s posttest. ****P<0.0001, ***P<0.001, **P<0.01, *P<0.05. **D)** Co-cultures of P14 (left graph) or adult (right graph) SI crypts with isolated adult (black) or P14 (blue) mesenchymal cells in factor-free medium. Whereas adult trophocytes support robust organoid growth, P14 trophocytes are inactive. Graph depicts organoids per sample relative to control wells (- cells + **E**GF, **N**OG, and **R**SPO1 (ENR); 94.5 ±46.8 P14 organoids/well; 64.5 ±20.03 adult organoids/well n=6 each). Statistical significance determined by Student’s t-test at **p<0.01, ***p<0.001 (n=3-6 biological replicates). **E)** P14 trophocytes support organoid growth in the presence of low concentrations of rEGF plus rNOG (EN medium) or rRSPO1 (ER medium) that are by themselves insufficient for organoid growth. Organoid counts are represented relative to parallel control crypt cultures in ENR medium without additional cells (59 ±12.1 organoids/well, n=4). Statistical differences determined by Student’s t-test at ***p<0.001 (n=4-6 biological replicates). See also Figure S1.

Newborn mouse ISCs, which reside in inter-villus troughs before crypt morphogenesis (Itzkovitz et al., 2012; Kim et al., 2012), and the organoids they generate differ from adult ISCs (Fordham et al., 2013). A watershed period between P10 and P16 is marked by crypt deepening prior to emergence of adult ISC properties and extensive crypt fission (Al-Nafussi and Wright, 1982; Cheng and Bjerknes, 1985; Langlands et al., 2016; Maskens and Dujardin-Loits, 1981; St Clair and Osborne, 1985; Sumigray et al., 2018a). To ask if PDGFRA^lo^ trophocytes, a principal BMPi source in adult mice (Greicius et al., 2018; Kim et al., 2020; McCarthy et al., 2020b; Stzepourginski et al., 2017), are active in this period, we co-cultured SI crypts with various PDGFRA^+^ cell types from P14 or adult *Pdgfra^H2B-eGFP^* mice. In contrast to adult trophocytes, which support organoid growth without any extrinsic factors, P14 trophocytes did not (Figures 2D and S1F). In the presence of rEGF plus low levels of rBMPi or rRSPO that did not alone support organoid growth, P14 trophocytes showed modest activity (Figure 2E). Thus, young trophocytes lack essential adult features and other stromal cells likely contribute ISC support, at least before weaning and possibly also in adults.

### Delineation of a multi-component sub-cryptal signaling center

To identify such cells, we extracted SI mesenchyme at various times between P1 and P14 for single-cell (sc) RNA-seq. We profiled >61,000 cells, encompassing unfractionated wild-type mesenchyme and GFP^+^ cells from *Pdgfra^H2B-eGFP^*mice (Supplemental Methods and Figure S2A). CD45^+^ (PTPRC^+^) leukocytes in some samples included macrophages, mast cells, and B and T lymphocytes, and early after birth we detected residual proliferation of Purkinje neurons, neural crest-derived cells, and trophocytes (Figure S2B-C). After excluding replicating cells, leukocytes, and Peyer’s patch-associated follicular cells (Link et al., 2007), >51,000 informative cells from all stages included all known adult populations (Figures 3A and S2D-E).

**Figure 3.**
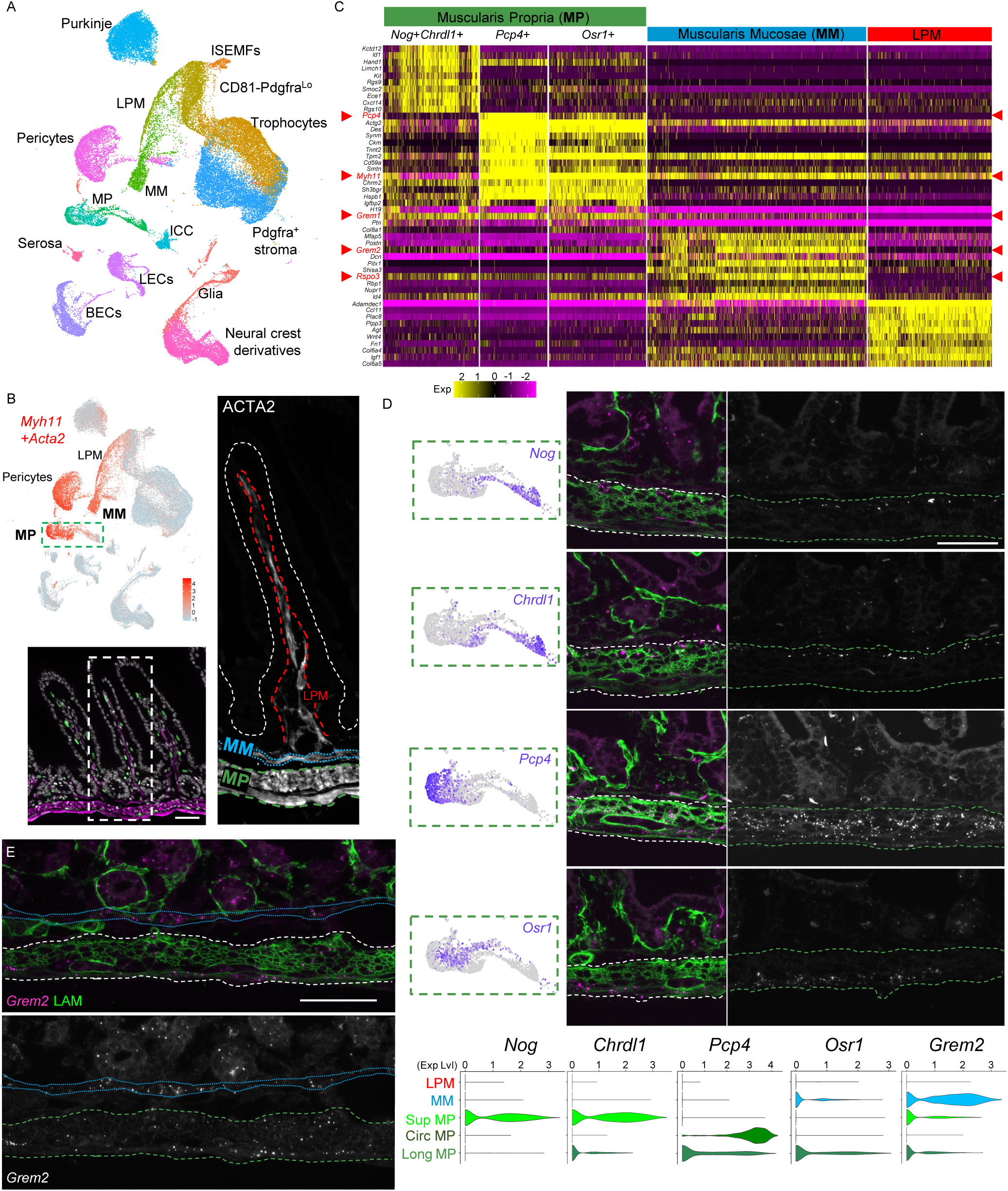
Diverse postnatal intestinal smooth muscle (SM) cell populations. **A)** Uniform manifold approximation and projection (UMAP) of scRNA-seq data from whole mesenchyme and GFP^+^ cells from *Pdgfra^H2B-eGFP^* mice, representing 51,084 cells from P1/P2, P4/P5, P9, and P14. MP, muscularis propria; LPM, lamina propria myocytes; MM, muscularis mucosae; ISEMFs, intestinal subepithelial myofibroblasts; ICC, interstitial cells of Cajal; LECs, lymphatic endothelial cells; BECs, blood endothelial cells. **B)** Overlay of aggregate *Myh11*+*Acta2* expression (red) on the UMAP plot, showing expression of these SM markers in each SM population and pericytes. ACTA2-immunostained P14 *Pdgfra^H2B-eGFP^* proximal SI (*Pdgfra^H2B-eGFP^* green, ACTA2 magenta, DAPI grey) highlights key anatomic structures magnified in the adjoining image (right – ACTA2 only): LPM (dashed red outline), MM (dotted blue outlines), and MP (dashed green outlines; epithelial surface in white dashed outline). Scale bar 50 μm. **C)** Top genes identified as discriminating markers for each indicated SM cell population. MP divides into *Nog*^+^*Chrdl1*^+^, *Pcp4^+^*, and *Osr1*^+^ subpopulations. Red arrowheads point to genes of particular interest; additional differences are listed in Table S2. **D)** RNAscope in situ hybridization (ISH) with the indicated probes (magenta dots on left, grey in right images, LAMININ=green). Three distinct MP layers (within dashed white or green outlines) correspond to scRNA-identified subpopulations (UMAP plot from the dashed green box in **B**): superficial *Nog^+^ Chrdl1^+^* cells, inner circular *Pcp4^+^* SM, and outer longitudinal *Osr1^+^* SM. Violin plots depict SM marker expression across indicated cell types. Sup, superficial; Circ, circular SM; Long, longitudinal SM. Scale bar 50 μm. **E)** RNAscope ISH for *Grem2* at P14 showing expression in MM (dotted blue outline) and MP (dashed green outline). Top, 2-color fluorescence; bottom, *Grem2* ISH in grey. Scale bar 50 μm. See also Figure S2 and S3.

Because adult SM, broadly defined, also expresses BMPi (Koppens et al., 2021; Martin-Alonso et al., 2021) and *Nog* is present in fetal SM fibers (Huycke et al., 2019), we first delineated specific BMPi sources within the *Acta2^hi^ Myh11^hi^* SM compartment. Graph-based cell clustering revealed molecular distinction among SM cells historically classified by their anatomic locations: lamina propria myocytes (LPM), muscularis mucosae (MM), and the muscularis propria (MP) comprised of inner circular and outer longitudinal SM layers (Figure 3B, note: pericytes also express high *Acta2* and *Myh11*). Whereas longitudinal and circular MP are largely similar, a novel MP-like population uniquely expresses the BMPi *Nog* and *Chrdl1* and low levels of *Acta2, Actg2,* and *Myh11* compared to other SM cells (Figure 3C and Table S2). In situ hybridization (ISH) of P14 intestines localized these unique *Nog^+^ Chrdl1^+^* cells to the superficial MP surface, closest to crypts and distinct from deeper *Pcp4^hi^* circular or *Osr1^+^*longitudinal MP cells (Figure 3D). These ISH distributions (quantitation and negative controls shown in Figure S3A-B) mirrored the marker specificity in scRNA data (Figure 3C-D).

Unlike MP, both LPM and MM express *Hhip*, as we verified by ISH (Figure S3C). Villus LPM is physically contiguous with sub-cryptal MM (Figure 3B), but the populations are molecularly distinct: whereas LPM lacks any BMPi and all other SM cells express *Grem1*, MM uniquely expresses high *Grem2* (Figures 3C-E and S3C-D). Double ISH with *Nog* and *Grem2* probes reliably distinguished *Grem2^+^* MM from *Nog*^+^ superficial MP (Figure S3E). Notably, each sub-cryptal population expresses *Rspo3* and at least two BMPi (Figure 4A). Taken together, this previously unappreciated SM cell and BMPi diversity near crypts reveals that, in addition to trophocytes, postnatal ISCs lie near tissue layers rich in key trophic factors.

**Figure 4:**
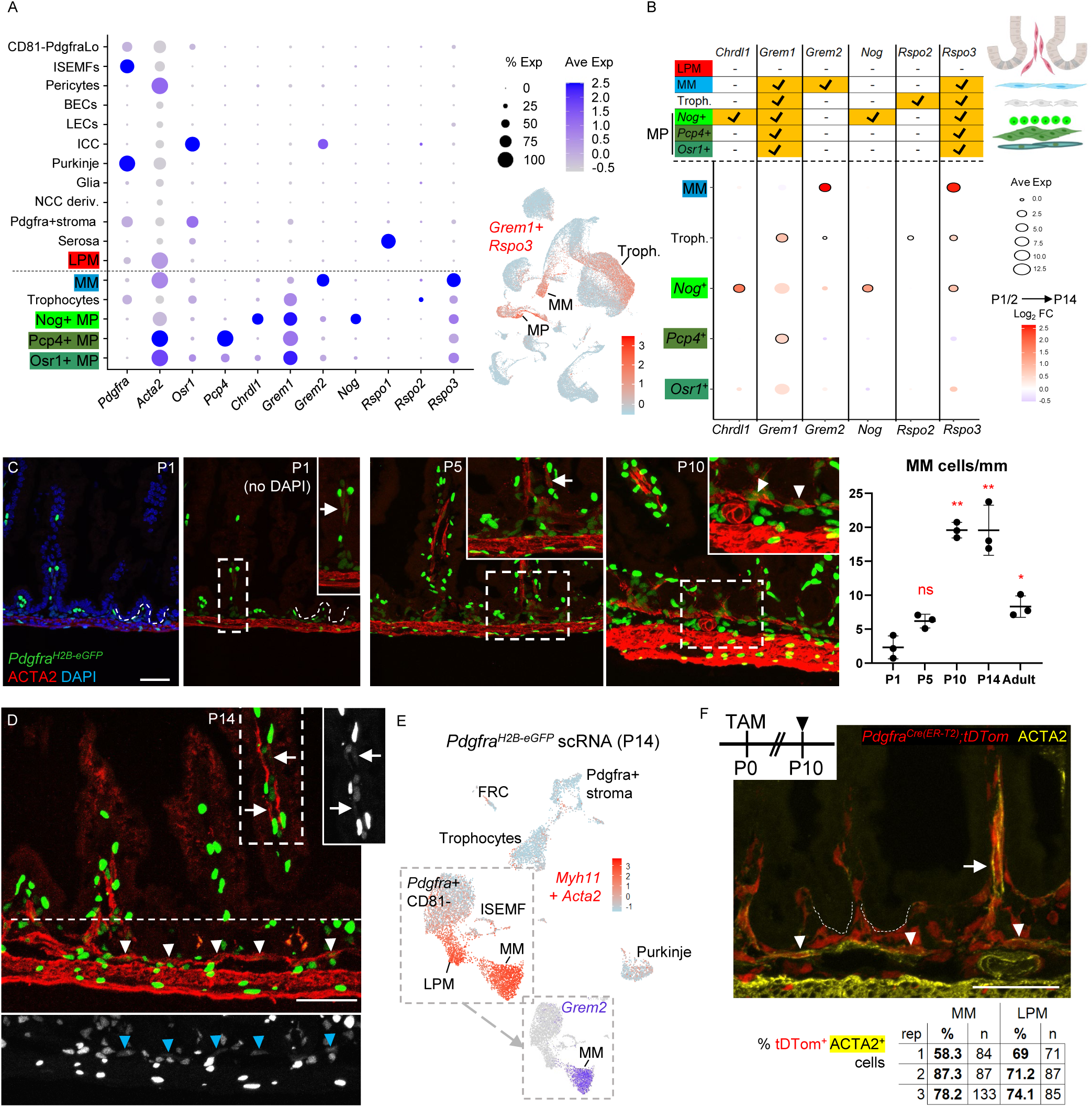
Sub-cryptal mesenchyme constitutes an ISC signaling center and the MM arises postnatally from native PDGFRA^+^ precursor cells. **A)** Average expression of selected BMPi and Rspo genes across all cell types represented in the postnatal mesenchymal scRNA survey (Figure 3A). Right, aggregate *Grem1* and *Rspo3* levels projected on the global UMAP show high dual expression in trophocytes (Troph.), MM, and MP. **B)** Top: schema for expression of key trophic factors in each distinct SM cell fraction in SI at P14. Bottom: log_2_ fold increase between P1/2 and P14 in average mRNA expression per cell of the indicated trophic factor genes. BMPi and Rspo gene expression increases significantly (*p* <0.05, black outline) in diverse postnatal sub-cryptal cells. **C)** Representative ACTA2 immunostaining of proximal *Pdgfra^H2B-eGFP^*SI at P1, P5, and P10. Boxed areas are magnified in insets. While LPM is present at P1 (arrow), MM arises between P5 and P10 (arrowheads). Scale bar 50 μm. Graph depicts MM cells/mm tissue at indicated ages. Statistical differences with respect to P1 were determined by one-way ANOVA followed by Dunnet’s multiple comparisons test. **p<0.0001, *p<0.05, n=3 animals at each age. **D)** GFP^+^ nuclei are evident in LPM (arrow) and MM (arrowheads) in *Pdgfra^H2B-eGFP^* proximal SI at P14, in agreement with scRNA data on purified GFP^+^ cells. Areas outlined with white dashes are shown to the right or below with GFP signals only in grey. Scale bar 50 μm. **E)** Aggregate *Acta2* and *Myh11* expression (red) overlaid on a composite UMAP plot of GFP^+^ cells isolated from *Pdgfra^H2B-eGFP^* mice at P14. FRC, fibroblastic reticular cells. Inset shows *Grem2* expression projected on the cells boxed in the composite UMAP plot. **F)** *Pdgfra^Cre(ER-T2)^;R26R^TdTom^* pups were treated with tamoxifen at P0 and their intestines were examined at P10, when LPM (arrow) and MM (arrowheads) showed *Pdgfra* lineage tracing. White dashes outline the epithelial-mesenchymal boundary. Scale bar 50 μm. tdtomato*^+^* MM and LPM cell fractions are shown (n: number of cells counted) from >4 mm of proximal SI in each of 3 independent animals. See also Figure S4 and S5.

### Adult niche structure represents the culmination of dynamic postnatal gene activity and de novo MM genesis

Other factors pertinent to niche functions include secreted Wnt antagonists in multiple cell types, canonical *Wnt2b* in trophocytes and MM and *Wnt2* in some lymphatic endothelial cells, and prominent BMP enrichment in ISEMFs, similar to adults (McCarthy et al., 2020b); various mesenchymal cells also express non-canonical *Wnt4* and *Wnt5a* (Figure S4A). Trophic factor levels increase in diverse sub-cryptal cells from birth to P14 (Figure 4B), together with those of tissue remodeling genes for example (Figure S4B). These postnatal mesenchymal dynamics may reflect in ISC supportive functions such as emergence of full trophocyte activity only after P14 (Figure 2D-E).

Our scRNA study recovered many more MM and LPM cells at P14 than earlier (Figure S2E). In line with this finding, ACTA immunostaining revealed sparse MM at P1, modest increase from P5 to P10, and a uniform structure consistently present only by P14 (Figure 4C-D). Whereas adult SM cells lack *Pdgfra* (or GFP in *Pdgfra^H2B-eGFP^*mice) (McCarthy et al., 2020b), large fractions of MM and LPM cells at P10 and P14 carried GFP^+^ nuclei (Figures 4C-D and S5A). Moreover, in our temporal scRNA survey, the 8,030 *Pdgfra^+^* (GFP^+^) cells from *Pdgfra^H2B-eGFP^*mice included distinct pools of *Acta2*^+^ *Myh11*^+^ cells corresponding to *Hhip^+^* LPM and *Grem2^+^ Rspo3^+^*MM (Figures 4E and S5B). Together, these findings suggest that MM and LPM derive from resident PDGFRA^+^ precursors, retaining H2B-eGFP (which is stable (Kanda et al., 1998)) for some duration. To test that hypothesis, we generated *Pdgfra^Cre(ER-T2)^*;*R26R^L-S-L-TdTom^*mice (Chung et al., 2018), (Madisen et al., 2010) (Table S1), induced Cre activity at P0, and followed the resulting TdTom label into *Pdgfra*^+^ cell derivatives at P10. PDGFRA^lo^ stroma and PDGFRA^hi^ ISEMFs were marked as expected, and LPM and MM also expressed TdTom (Figures 4F and S5C). Thus, these SM cells arise *de novo* from resident PDGFRA^+^ stromal cells well after birth.

In adult mice, MM is anatomically distinct from the MP (Figure S5D) and, compatible with niche activity, its *Grem2^+^* cells lie directly beneath *Olfm4^hi^* ISCs at the crypt base (Figure 5A). To determine if MM retains its key early postnatal products, we generated *Myh11^Cre(ER-T2)^;Pdgfra^H2B-eGFP^*;*R26R^TdTom^* mice, where SM cells label red and PDGFRA^+^ cells green (Figure 5B and Table S1). After manually stripping the MP, scRNA analysis of TdTom^+^ GFP^-^ cells from the SI yielded few *Grem2^+^* MM cells (Figure S5E); integration with previous data from MP-depleted adult SI mesenchyme (McCarthy et al., 2020b) revealed *Grem2*^+^ *Rspo3*^+^ MM distinct from LPM (Figure S5F). Because MM is thicker in the colon than in the SI (Figure 5C), we anticipated a higher cell yield from adult colon and, after stripping colonic MP (including superficial *Nog^+^ Chrdl1^+^* cells) scRNA-seq of TdTom^+^ GFP^-^ cells identified a distinct *Grem1^+^ Grem2^+^ Rspo3^+^* cell population corresponding to MM, separate from abundant *Hhip^+^* LPM (Figure 5C). ISH for these markers in adult colon (Figure 5D) and SI (Figure S6A-C) verified the expression domains that scRNA-seq had revealed in sub-cryptal MM, trophocytes, and superficial MP.

**Figure 5.**
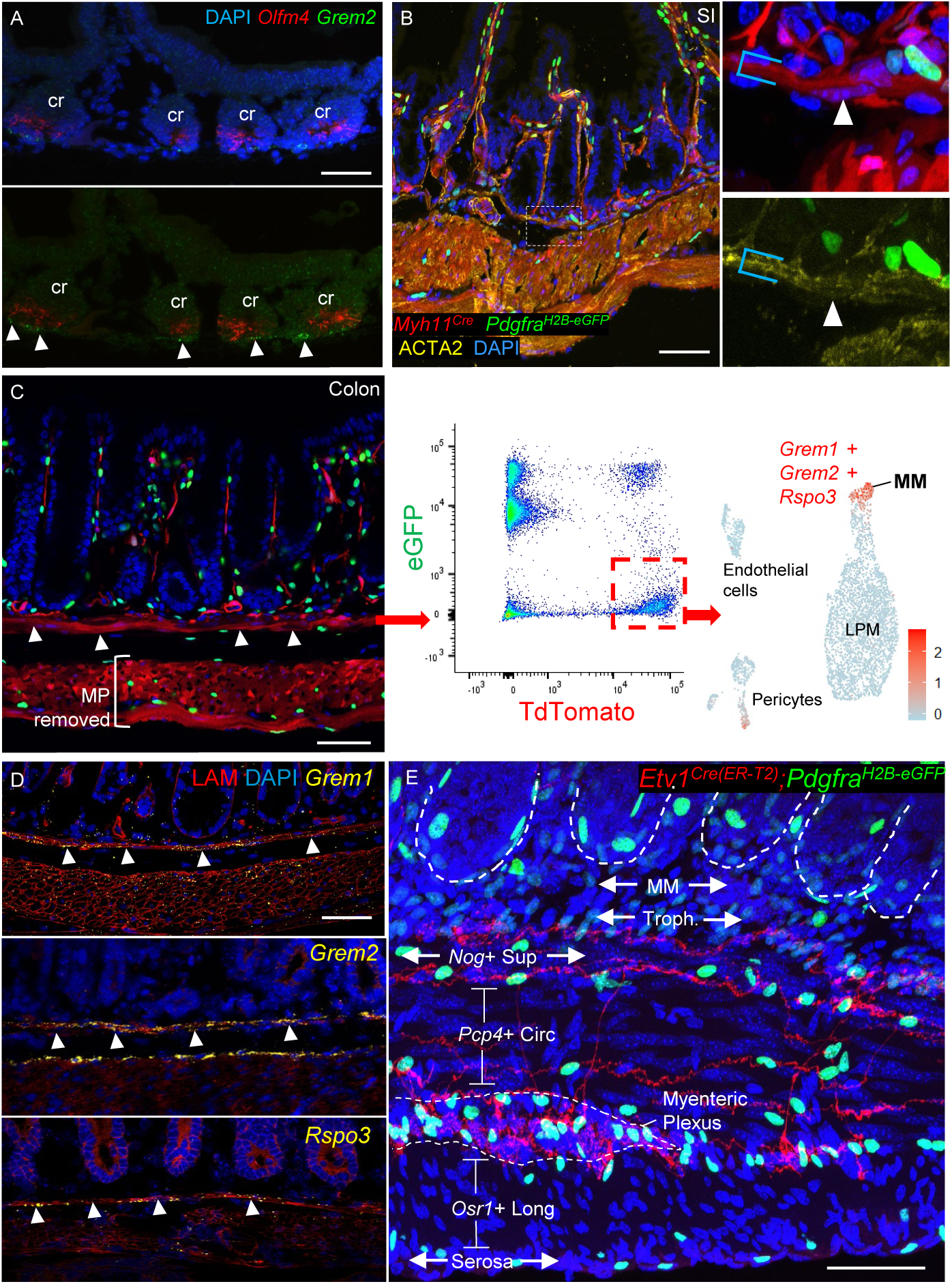
Sub-epithelial cells together constitute an ISC signaling center in adult mice. **A)** *Grem2* expression revealed by RNAscope ISH (green) in adult duodenal MM, directly beneath *Olfm4^+^* (red) ISCs (arrowheads); cr, crypts. MP was removed manually before staining and the bottom image is the same as the top, with DAPI signal excluded. Scale bar 50 μm. **B)** Adult *Myh11^Cre(ER-T2)^;R26R^TdTom^;Pdgfra^H2B-eGFP^* SI showing GFP^+^ Tom^+^ ISEMFs and Tom^+^ SM cells. The dashed box is magnified in the right images, where the bracket outlines MM and the arrowhead points to a GFP^-^ cell characteristic of adult MM. Scale bar 50 μm. **C)** Adult *Myh11^Cre(ER-T2)^;R26R^TdTom^;Pdgfra^H2B-eGFP^* colon showing thick MM (arrowheads). The MP was removed prior to FACS sorting and a representative FACS plot shows colonic Tom^+^ GFP^-^ cells (dashed red box), which were isolated for scRNA analysis. A UMAP plot of 3,592 cells shows aggregate *Grem1*, *Grem2*, and *Rspo3* expression in the small MM fraction, distinct from LPM, endothelial cells or pericytes. Scale bar 50 μm. **D)** RNAscope ISH for *Grem1*, *Grem2*, and *Rspo3* (all signals in yellow dots, red: LAMININ) in adult mouse colon, showing predominant expression in MM (arrowheads) and superficial MP. **E)** Representative whole-mount image from *Etv1^Cre(ER-T2)^;R26R^TdTom^;Pdgfra^H2B-eGFP^* mouse ileum, showing the tissue beneath crypts (dashed white outlines) in relation to axonal projections (red) from *Etv1^+^*interstitial cells of Cajal that demarcate SM layers. *Pdgfra^lo^*trophocytes lie between sub-cryptal MM (unstained) and *Nog^+^ Chrdl1^+^* superficial MP (also unstained; neural projections separate these cells from circular *Pcp^+^* SM). GFP^hi^ nuclei far from peri-cryptal ISEMFs are Purkinje neurons (Kurahashi et al., 2012b), distinct from *Etv1^Cre^*-labeled TdTom^+^ projections. Scale bar 50 μm. See also Figures S5 and S6.

Adult *Pdgfra^lo^Grem1^+^* trophocytes consistently localize between *Grem2*^+^ MM and superficial *Nog^+^* MP. To resolve closely apposed mesenchymal cell layers at fine resolution, we generated *Etv1^Cre(ER-T2)^;R26R^TdTom^;Pdgfra^H2B-eGFP^*mice, where Cre activity marks axons projecting from ETV1^+^ interstitial cells of Cajal (ICC) (Chi et al., 2010) (Table S1). PDGFRA^lo^ trophocytes are indeed sandwiched between the MM and superficial MP (Figure 5E, note: intra-muscular PDGFRA^hi^ cells are Purkinje neurons (Kurahashi et al., 2012a)). Altogether, these findings reveal diverse SM populations that flank trophocytes, mature in concert with organ and crypt growth, and by virtue of BMPi and RSPO expression likely constitute a layered signaling hub.

### SM requirements for ISC support in vivo

Duplicative and wide expression of niche factors in distinctive sub-cryptal cells hints at their likely joint and redundant contributions to an ISC signaling center. Ablation of adult *Grem1^+^* cells triggers epithelial BMP activity and rapid ISC attrition (McCarthy et al., 2020b) but because many mesenchymal cells express *Grem1* (Figure 4A), this finding does not implicate a defined single niche component. To isolate SM contributions to the ISC niche *in vivo*, we generated *Myh11^Cre-ER(T2)^*;*R26R^L-S-L-DTA^*;*Pdgfra^H2B-eGFP^*mice (*Myh11;DTA*, Table S1); Cre activity in *Myh11^Cre-ER(T2)^*;*R26R^L-S-L-tDtomato^*mice mirrors ACTA2 immunostaining (Figure S6D). Adult *Myh11;DTA* mice became moribund within 2 days of tamoxifen (TAM) exposure, likely as a result of global SM attrition and precluding assessment of intestinal niche functions. However, administering TAM at P14, by when the MM is well formed, allowed us to follow animals for 1 week. Weight loss and proportionally reduced SI length indicated muscle attrition (Figure S7A). ACTA2^+^ MM was substantially depleted at P16 and P21 (Figure 6A); circular MP was modestly reduced at P21 (Figure S7B); PDGFRA^hi^ ISEMFs were preserved; and PDGFRA^lo^ cells were slightly reduced, likely reflecting loss of PDGFRA^+^ SM precursors (Figure S7C). Thus, SM ablation was incomplete in the study timeframe, as reported with *R26R^DTA^* and other Cre drivers (Eilken et al., 2017) or owing to limited TAM activity in intestinal sub-epithelium (Chee et al., 2018). Nevertheless, MM appeared especially vulnerable and TAM-treated *Myh11;DTA* pups were informative with respect to SM functions in the pre-weaning ISC niche.

**Figure 6.**
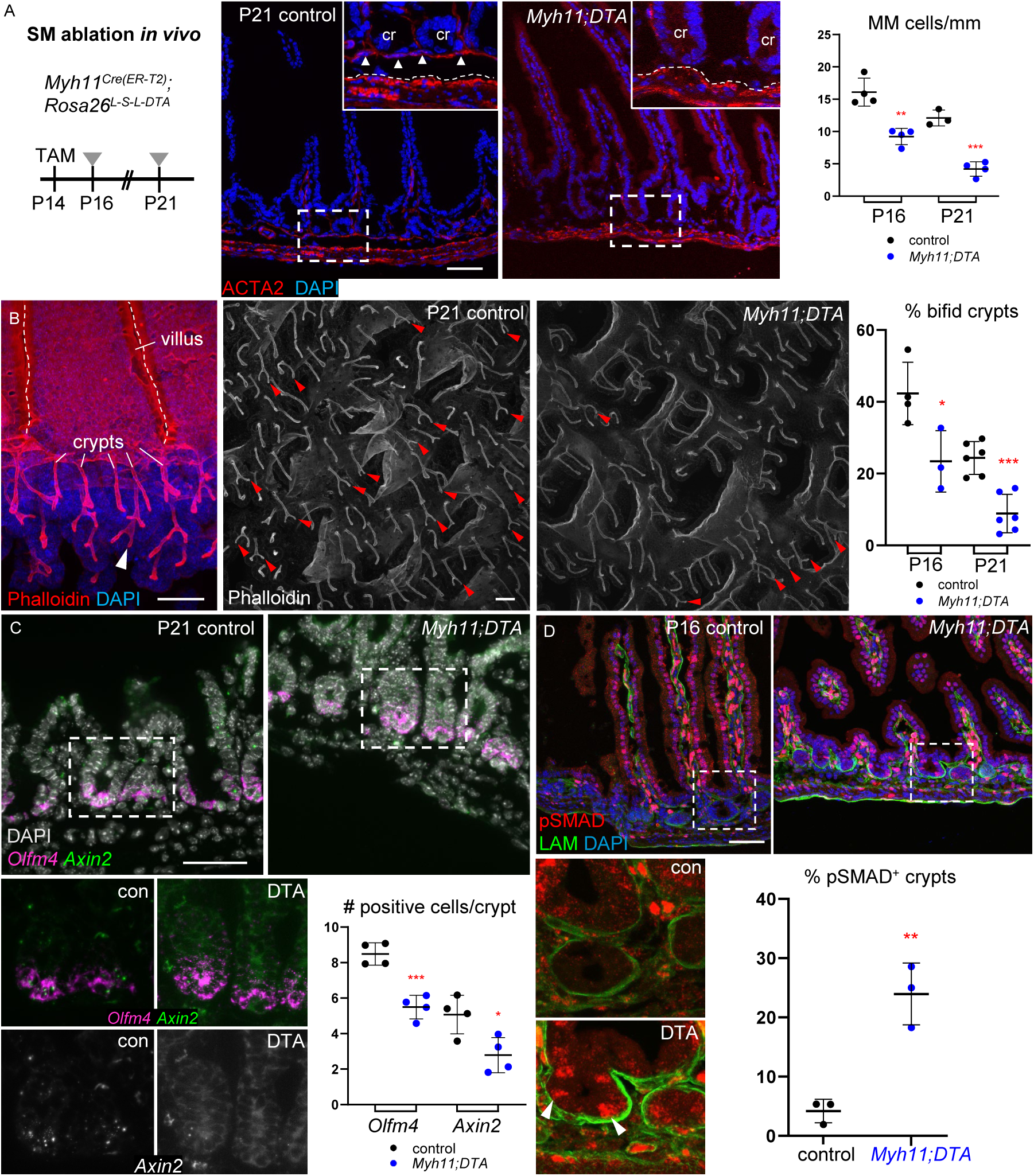
BMP-associated crypt attrition in mice with intestinal SM deficiency. **A)** *Myh11^Cre(ER-T2)^;R26R^DTA^* mice were treated with tamoxifen at P14 and intestines harvested at P16 and P21 reveal reduced ACTA2-stained MM. Representative images at P21 are shown and boxed areas are magnified in the insets, where the bracket demarcates control MM and dashed white lines mark the superficial MP border; cr: crypts. Scale bar 50 μm. To the right, MM cells are quantified from >4 mm tissue in each of 3 or 4 independent animals. Statistical significance was determined using unpaired Student’s t-test. **p<0.01, ***p<0.001. **B)** F-actin (phalloidin)-stained epithelial whole-mount images showing considerably fewer bifid crypts (arrowheads) in *Myh11^Cre(ER-T2)^;R26R^DTA^* compared to control pups injected with tamoxifen (TAM) at P14 and examined at P16 (Figure S7A) and P21. To the right, bifid crypts are quantified from >100 crypts in each of 3-6 mice from each age. Statistical significance was determined using unpaired Student’s t-test. *p <0.05, ***p<0.001. **C)** Crypt base cells expressing RNA for ISC markers *Olfm4* (RNAscope ISH signal in magenta) and *Axin2* (FITC) are decreased in TAM-treated *Myh11;DTA* animals at P21. n=4 mice per condition, >40 crypts per sample, statistics by unpaired Student’s t-test. *p<0.05, ***p<0.001. **D)** Increased pSMAD1/5 immunostaining in TAM-treated *Myh11^Cre(ER-T2)^;R26R^DTA^* crypts (arrowheads), which lack pSMAD1/5 in controls (see also Figure 1A). pSMAD1/5^+^ crypts were quantified in >80 crypts from 3 mice for each condition and statistical significance determined using unpaired Student’s t-test. **p<0.01. See also Figure S7.

Crypt cell proliferation, which occurs largely in transit amplifying cells, was intact (Figure S7D) but crypt fission, which peaks in the 3^rd^ week of mouse life (Al-Nafussi and Wright, 1982), was significantly attenuated by P21 and starting as early as P16 (Figures 6B and S7E). ISC markers *Olfm4* and *Axin2* were also reduced at P21 (Figure 6C) and crypts with pSMAD^+^ cells were increased ∼5-fold (Figure 6D), pointing to BMPi deficiency as a basis for these defects. Thus, SM deficiency compromises ISC function, owing at least in part to unopposed BMP signaling.

### Complementary ISC support from SM and trophocytes

*Myh11;DTA* mice implicate SM –but not a discrete SM cell type– in ISC support, and because the above findings do not address the relative contributions of trophocytes and SM, we assessed SM niche activity *in vitro*. A lack of selective surface markers precludes purification of MM or *Nog*^+^ superficial MP by flow cytometry. First, we therefore cultured unfractionated P14 or adult MP with P14 or adult SI crypts. Neither young nor adult whole MP substituted for ENR medium in supporting organoid formation; however, in the presence of low doses of rEGF and rRSPO1, which were insufficient for organoid growth, MP co-cultures consistently generated organoids (Figures 7A and S7F). Thus, although MP lacks the potency of trophocytes in this assay, its activity is sufficient to spawn organoids when RSPO levels are limiting.

**Figure 7.**
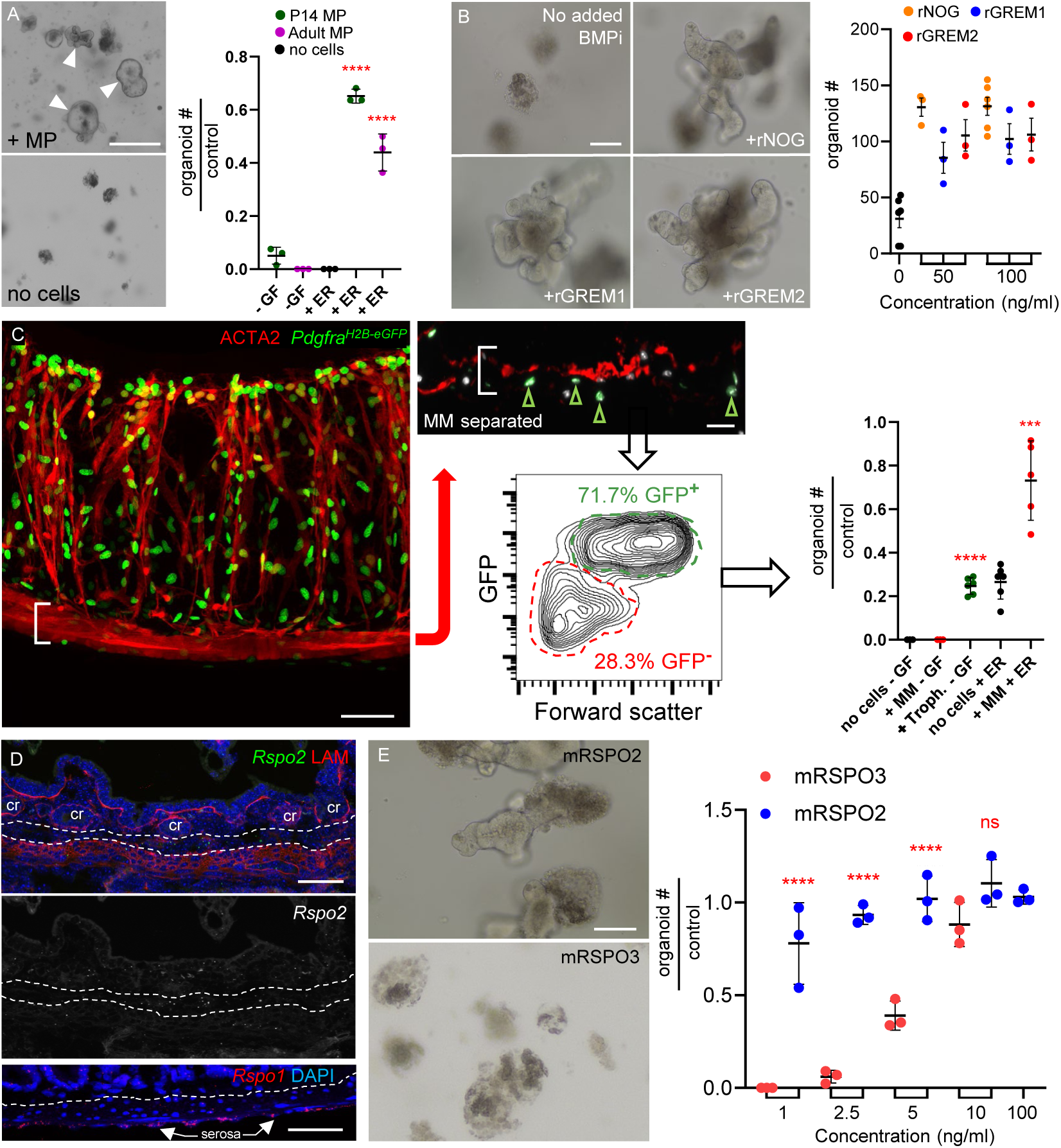
Adult MM and MP complement trophocyte activity *in vitro* by substituting for BMPi. **A)** Co-culture of adult SI crypts with unfractionated P14 or adult MP did not generate organoids (arrowheads) by itself but did so in the presence of sub-optimal concentrations of rEGF and rRSPO. Organoids are represented in the graph relative to the numbers generated in ENR medium (no added cells, 66 ±10.4 organoids, n=3). Significance determined by Student’s t-test, ****p<0.0001. -GF, no recombinant trophic factors. **B)** Organoids arising from SI crypts after 5 days of culture in a sub-optimal concentration of rRSPO1 (see Methods) and 50 ng/mL or 100 ng/mL of the indicated BMP inhibitors. n=3 biological replicates. One-way ANOVA with Dunnett’s test shows significance of all conditions to at least p<0.01 compared to controls with no added BMPi. **C)** After MP removal, MM stripped from *Pdgfra^H2B-eGFP^* colon retained GFP^+^ trophocytes on the undersurface (green arrowheads in inset) and flow cytometry separated GFP^+^ trophocytes from GFP^-^ MM (see Figure S7G-H). Scale bars 50 μm. In adult SI crypt co-cultures, GFP^+^ cells supported organoid growth without added factors, while GFP^-^ MM provided support only in the presence of inactive amounts of rEGF and rRSPO1 (ER medium). Significance determined with respect to ENR-only controls (no cells, 31.7 ±5.5 organoids/well, n=5) using Student’s t-test. ***p<0.001, ****p<0.0001. **D)** Top: *Rspo2* (green) ISH and LAMININ (red) immunostaining in adult SI. Dashed lines indicate boundaries marked by the MM and superficial MP, between which lie submucosal trophocytes (cr: crypts). Greyscale version of the *Rspo2* signal is shown below the color image without LAM. Bottom: *Rspo1* (red) ISH confirms high expression in the serosa as indicated by scRNA-seq. Dashed line marks superficial MP. Scale bars 50 μm. **E)** Crypts cultured with as little as 1 ng/mL rRSPO2 form organoids at an efficiency that requires at least 10 ng/mL rRSPO3. Graph depicts organoid-forming efficiency at 5 days relative to 100 ng/ml rRSPO3 (107.3 ±19.6 organoids/well, n=3). Statistical comparisons used one-way ANOVA followed by Tukey’s posttest. ****P <0.0001, scale bar 50 μm. See also Figure S7.

Second, noting that MM is an exclusive source of *Grem2*, we found that rGREM2 supported organoid growth as potently as rGREM1 (Figure 7B), indicating that MM is also well situated to complement trophocyte activity. Indeed, MM isolated physically as a membrane from adult *Pdgfra^H2B-eGFP^*colon supported organoid growth from adult SI crypts in the absence of additional factors (Figure S7G); this effect could, however, reflect trophocyte contamination. We therefore digested the intact membrane, sorted GFP^-^ (MM) from GFP^+^ (trophocyte) cells by flow cytometry (Figure 7C), and verified that GFP^+^ cells are CD81^+^ trophocytes (Figure S7H). The cell yields allowed us to co-culture only 10,000 cells from each fraction with SI crypts in small Matrigel droplets, with limited flexibility in factor supplementation; nevertheless, the trophocyte fraction promoted organoid growth in the absence of soluble factors. Although MM was inert by itself, in the presence of small amounts of rEGF and rRSPO1, which approximated trophocyte activity in this assay, its supportive activity consistently exceeded that of trophocytes alone (Figure 7C). Thus, MP and MM each reveal intrinsic and complementary niche activities when factor levels are sub-optimal, and organoid growth in the absence of rNOG suggests that sub-cryptal SM is a functional source of BMPi.

Finally, because adult trophocytes alone can expand ISCs robustly *in vitro* (McCarthy et al., 2020b) (see Figs. 2D and 7C), these cells must harbor some distinctive activity, not limited to BMPi. We noted that several cell types express *Rspo3* and serosal cells express *Rspo1* (*Rspo4* is absent from any cell type), but only trophocytes express *Rspo2* (Figure 4A); we confirmed this finding by ISH (Figure 7D). RSPO3 is superior to RSPO1 in sustaining organoids (Greicius et al., 2018) and ISC compromise *in vivo* requires both RSPO2 and RSPO3 antibodies (Storm et al., 2016), but RSPO2 activity has not been examined in isolation. We found that adult mouse SI crypts required 10 ng/mL murine rRSPO3 to generate organoids at top efficiency and 2.5 ng/mL was ineffective, but as little as 1 ng/mL rRSPO2 achieved near-peak efficiency (Figure 7E) and human RSPO2 also was more potent than RSPO1 or RSPO3 (Figure S7I). Thus, redundant BMPi coupled with unique expression of highly potent RSPO2 likely explains *ex vivo* organoid support from trophocytes alone.

## Discussion

Immature mouse ISC precursors located in flat inter-villus epithelium first generate shallow troughs, which deepen into the earliest crypts and later undergo extensive fission to expand crypt and epithelial mass before weaning age (Al-Nafussi and Wright, 1982; Itzkovitz et al., 2012; Kim et al., 2012; Schmidt et al., 1988; Sumigray et al., 2018a). Classical (Al-Nafussi and Wright, 1982; Bry et al., 1994; Cheng and Bjerknes, 1985; St Clair and Osborne, 1985) and recent (Sumigray et al., 2018a) studies highlight the P10-P15 interval as a watershed, when crypts and ISCs acquire adult form and cell composition. Our study integrates mesenchymal scRNA profiles with high-resolution anatomic clarity and functional assessment of the emerging ISC niche during this watershed period. ISEMF aggregation at the villus isthmus coincides with elevated BMP signaling in adjacent epithelial cells and, likely in part to counter that increase, sub-epithelial niche components develop in parallel. One previously unknown *Acta2^lo^ Myh11^lo^* population within superficial circular MP fibers uniquely expresses the BMPi *Nog* and *Chrdl1*, and mRNA expression of multiple trophic factors increases postnatally in these cells and in trophocytes. Sub-cryptal MM, a previously unappreciated niche component expressing *Rspo3* and BMPi, is distinct in this respect from perpendicular and contiguous LPM fibers and it arises *de novo* after birth from resident PDGFRA^+^ mesenchyme. Altogether, these findings identify a finely layered hub of partially redundant RSPO- and BMPi-expressing cells that position strategically near the crypt base to fulfill adult epithelial needs.

Previous ablation of *Grem1^+^* cells, likely including SM as well as trophocytes, affected ISCs profoundly (McCarthy et al., 2020b). Selective depletion of the *Myh11^+^* SM compartment in the present study also precipitated ISC dysfunction, reflected in markedly reduced crypt fission that normally peaks in the third week of mouse life. Given that P14 trophocytes fail to support organoids without added BMPi, these findings indicate that overlapping activities of cells with overlapping expression of trophic factors help drive postnatal crypt expansion and subsequent ISC support. For example, postnatal emergence of MM may specifically counteract ISEMF aggregation at the villus base. Adult MM and MP both harbor discrete niche activities, providing BMPi but likely insufficient RSPO for organoid growth.

Our findings add developmental and fine spatial context to prior work showing BMPi activity in driving ISC properties (Batts et al., 2006; Davis et al., 2015; Haramis et al., 2004; He et al., 2004), polarized BMP and BMPi expression in human colonic mucosa (Kosinski et al., 2007), and that BMP signaling restrains both ISC pool size (Qi et al., 2017) and crypt fission (Auclair et al., 2007) independent of Wnt signaling. Furthermore, biochemical (Yan et al., 2017), functional (Storm et al., 2016), and structural studies (Chen et al., 2013; Peng et al., 2013; Wang et al., 2013; Zebisch et al., 2013) point to sequential Wnt and RSPO activity within the same signaling pathway, with canonical Wnts priming ISCs to express RSPO receptors. In addition to one or more BMPi, every sub-cryptal mouse niche component expresses at least one *Rspo* gene and trophocytes’ unique expression of the most potent member, *Rspo2*, may explain their particularly robust support of organoid growth (McCarthy et al., 2020b).

## Limitations of the study

Establishing the functions and requirements of each niche component *in vivo* is a tall order. Isolation or ablation of specific cell populations requires selective surface markers or Cre-driver mouse strains that may target multiple cell types. *Myh11^Cre^*, for example, marks all SM cells, including LPM, MM and MP, hence limiting interpretation of phenotypes. Moreover, SM ablation was unavoidably incomplete and our findings in *Myh11^Cre^;DTA* mice could reflect indirect effects of SM depletion on other niche cells rather than direct consequences on nearby ISCs. Finally, although crypt fission is known to require opposition of BMP signaling (Auclair et al., 2007) and we observed increased pSMAD in *Myh11^Cre^;DTA* mice, it is arguably unclear whether BMPi or other SM products enable optimal crypt fission *in vivo*. Our systematic effort to delineate niche-element functions in organoid assays (Figure 7) substantially mitigates some of these limitations. In the future, molecular markers that allow further cell fractionation and precise *in vivo* ablation will complete the emerging understanding of individual niche elements. Of particular interest is the novel *Acta2^lo^ Myh11^lo^ Nog^+^ Chdl1^+^* cell layer, which should be characterized further. Combinations of the molecular markers identified in this study will aid in these endeavors.

## Acknowledgments

Supported in part by National Institutes of Health awards U01DK103152 and R01DK121540 (to R.A.S.) and K01DK125639 (to N.M.). We acknowledge generous research support from the Lind family; core microscopy and organoid culture services from the Harvard Digestive Diseases Center (P30 DK034854); and valuable assistance from E. Manieri and flow cytometry services at the Dana-Farber Cancer Institute.

## Author Contributions

N.M. and R.A.S. conceived and designed the studies; N.M. performed most experiments and analyses, with help from G.T. and A.M. for crypt co-cultures and S.M. for scRNA-seq analysis; J.K. performed ISH on colonic tissue; N.M. and R.A.S. drafted the manuscript, with input from all authors.

## METHODS

### EXPERIMENTAL MODEL AND SUBJECT DETAILS

#### Animals

*Pdgfra^H2B-eGFP^* (Jackson Laboratories (JAX) strain 007669) (Hamilton et al., 2003), *Rosa26R^LSL-TdTomato^* (JAX strain 007909), *Myh11^Cre(ER-T2)^* (JAX strain 019079), *Pdgfra^Cre(ER-T2)^*(JAX strain 032770), *Etv1^Cre(ER-T2)^* (JAX strain 013048) and *Rosa26R^LSL-DTA^* (JAX strain 009669) mouse lines were purchased from Jackson Laboratories (Table S1). Adult mice were more than 8 weeks of age at the time of treatments or cell isolations. All experiments used mice of both sexes and littermates as controls. All animal procedures and experiments were approved and monitored by the Animal Care and Use Committee at the Dana-Farber Cancer Institute.

### METHOD DETAILS

#### Mouse treatments

Postnatal *Pdgfra^Cre(ER-T2)^*;*R26R^LSL-TdTom^*and *Myh11^Cre(ERT2)^;Rosa26 ^LSL-DTA^* mice received 1 dose of 4-OH tamoxifen (Sigma-Aldrich, 1 mg per 25 g body weight) by gastric gavage at P1 or intra-peritoneal (IP) injection at P14. *Myh11^Cre(ER-T2)^;R26R^LSL-TdTom^;Pdgfra^H2B-eGFP^*adults received 4-OH tamoxifen (1 mg) by IP injection on 2 consecutive days to allow recombination at *LoxP* sites and were harvested at the indicated times, usually 5 days later. BrDU (10 μg/g body weight) was administered by IP injection 1 h before euthanasia. Postnatal mice were harvested at the time points indicated in the figure legends.

#### Immunohistochemistry and quantitation

Whole-mount tissue immunohistochemistry was performed as described (Bernier-Latmani and Petrova, 2016; McCarthy et al., 2020b). Briefly, proximal small intestines were harvested, pinned onto agarose plates, and fixed overnight in 4% paraformaldehyde (PFA); in this and all subsequent steps, the tissue was rocked gently. After rinsing in phosphate-buffered saline (PBS), the tissue was placed in 10%, then 20% sucrose over the course of 1 day, followed by blocking buffer (PBS containing 0.125% bovine serum albumin, 0.003% Triton X-100, 0.05% donkey serum, and 0.0005% NaN_3_) for 6 h and overnight in blocking buffer containing 4′,6-diamidino-2-phenylindole (DAPI). Tissue was rinsed with PBS, cut into 1-mm fragments, placed on glass slides with spacers (Grace Bio-Labs, 654002), and cleared using FocusClear (CelExplorer Labs, FC-101) for 30 min, before applying VectaShield mounting medium (Vector Laboratories) and a coverslip. To generate representative and comprehensive anatomic resolution, images were taken from at least 3 independent animals.

Proximal SI epithelial immunochemistry was performed as described (Sumigray et al., 2018b). Fresh tissue was incubated in 5 mM EDTA in Hank’s Balanced Salt Solution (HBSS) at room temperature, rocked gently for 5 min, and washed briefly in PBS before separating the epithelium gently from underlying mesenchyme. Tissue was fixed in 4% PFA at 4°C overnight, with subsequent PBS washing and incubation with DAPI to stain nuclei and Phalloidin (Invitrogen, A12381) to visualize filamentous actin. Crypt bifurcation was quantified in 3D-rendered images of whole-mount phalloidin stained intestines, reported as a fraction of >100 intact crypts.

Routine immunohistochemistry was performed on tissues fixed as described above and placed in OCT compound (Tissue-Tek, 4583). 7 μm sections were prepared using a Leica CM3050 cryostat. The following antibodies (Ab, all at 1:1000 dilution unless indicated) were used: Laminin (Sigma, L9393); GFP (Abcam, ab6662); CD31 (BD Biosciences, 557355); bromodeoxyuridine (BrDU, Life Technologies, B23151); Alpha-smooth muscle actin (ACTA2; Abcam, ab5694); PDGFRA (R&D Systems, AF1062, 1:100); and Alexa Fluor-conjugated secondary goat anti-rat, goat anti-rabbit, or donkey anti-goat IgG (Invitrogen, A11081, A21071, A11058). Signals were amplified using biotin/streptavidin HRP-conjugated secondary Ab (Jackson Immuno, 111-065-003 and 016-030-084) and detected using Tyramide Signal Amplification Plus kit (Akoya Biosciences, NEL744001KT, 1:100). Images were taken using a Leica SP5X laser scanning confocal microscope and further processed using ImageJ Fiji software (Schindelin et al., 2012).

pSMAD1/5 was immunostained as described (Nerurkar et al., 2017). Briefly, antigens were retrieved by boiling slides in citrate buffer (pH 6.0) for 1 min and pSMAD1/5 Ab (Cell Signaling, 41D10) was added overnight at 4°C. pSMAD1/5^+^ crypts were quantified on images acquired and processed as above. Epithelial compartments were delineated with respect to LAM Ab-stained basement membrane, and quantified by representing the lowest and highest 5 villus epithelial cells (10 cells per villus) as the fraction of pSMAD1/5^+^ cells in >25 villi per sample (Figure 1A) or crypts carrying ≥1 pSMAD1/5^+^ cell as a fraction of >80 crypts per sample (Figure 6D). To quantify *Pdgfra^H2B-eGFP^*ISEMFs, 25 μm sections were co-stained with LAM Ab to demarcate the epithelium. Mesenchymal GFP^hi^ cells abutting the bottom 5 villus epithelial cells were counted and are reported with respect to GFP^hi^ cells abutting the next 5 epithelial cells along the villus trunk (Figure 1C). More than 25 villi were counted per sample.

To quantify SM populations (Figures 6A and S7B), ACTA2^+^ cells in the MM and in the circular and longitudinal layers of the MP were counted per length of proximal SI examined (average 5 mm/sample). Co-labeled ACTA2^+^*Pdgfra^lo^* cells are reported as a ratio of the total number of ACTA2^+^ MM and LPM (Figures 4C and S5A). *Pdgfra^Cre(ER-T2)^;Rosa26^DTA^*intestines exposed to TAM at P0 were harvested at P10 and stained with ACTA2 Ab to quantify various SM compartments. For quantitation of *Pdgfra^lo^* cells in *Myh11^Cre^;Pdgfra^H2B-eGFP^;DTA* experiments (Figure S7C), cells found in the submucosa, between crypts and MP, are reported as numbers/length of intestine (>4mm/sample). BrdU+ cells/crypt are reported as numbers per crypt (≥25 crypts/sample).

#### In situ RNA hybridization and quantitation

mRNAs were localized by the RNAscope (Advanced Cell Diagnostics) method (Wang et al., 2012) on intestines collected from at least 3 different animals per treatment. Probe sets were designed by Advanced Cell Diagnostics for *Chrdl1*, *Grem1*, *Grem2*, *Hhip*, *Noggin*, *Olfm4*, *Axin2*, *Osr1*, *Pcp4*, *Rspo1*, *Rspo2*, and *Rspo3*. After hybridization according to the manufacturer’s protocols, tissue sections were washed for 5 min in PBS containing 0.1% Tween-20, blocked for 1 h at room temperature in PBS containing 5% normal goat serum, and exposed overnight at 4°C to Laminin (Sigma, L9393, 1:1,000) or GFP (Abcam, ab6556, 1:100) Ab. After multiple 5-min washes in PBS and 90-min incubation with AlexaFluor-conjugated secondary Ab as above (Invitrogen) at room temperature, DAPI was applied and slides were mounted according to the RNAscope protocol. Images were taken using a Leica SP5X laser scanning confocal or a Leica Thunder Imager microscope and processed using ImageJ Fiji software (Schindelin et al., 2012). For quantitation in SM layers delineated by laminin (Figure S3B and S3D, 2 samples), each cell with at least 1 fluorescent ISH dot was counted as one and reported as a fraction of all SM cells present in the respective sub-compartment. Every cell with at least one *Olfm4*+ or *Axin2*+ ISH dot was reported per crypt (Figure 6C, >40 crypts per sample).

**Schematic illustrations** were generated with BioRender.

#### Mesenchymal cell isolation and flow cytometry

Mesenchymal cells were isolated from the pooled proximal halves of the small intestine from 3-6 wildtype or *Pdgfra^H2B-eGFP^*pups. Timepoints harvested from wildtype tissue include P1, P4, P5, P9, and P14, and from GFP+ isolated *Pdgfra^H2B-eGFP^* tissue include P2, P5, and P14. After manual stripping of external muscles (MP) and serosa, whole adult *Myh11^Cre(ER-T2)^;R26R^L-L-TdTom^; Pdgfra^H2B-eGFP^* SI or colon was processed as described(McCarthy et al., 2020b). Epithelium was denuded by shaking the tissue for 20 min at 37°C in pre-warmed HBSS (Life Technologies) containing 10 mM EDTA. The remaining tissue was rinsed with HBSS, minced using a scalpel, and digested with gentle rocking for 1 h at 37°C in 3 mg/mL collagenase II (Worthington, LS004176) diluted in HBSS containing 5% fetal bovine serum (FBS). Extracted cells were centrifuged at 300 *g* for 5 min, washed with FACS buffer (PBS containing 0.1% BSA), and leukocytes were depleted in ACK Lysis buffer (Gibco) for 3 min. Washed cells were suspended in FACS buffer and, to deplete epithelial and immune populations, stained with conjugated EPCAM (BioLegends, 118214, 1:100) and CD45 (eBiosciences, 17-0451-82, 1:100) Ab for 20 min at 4°C. Cells were sorted on a FACSAria IIII flow cytometer, with gating against DAPI (BD Pharmingen) to identify live cells. Graphs of isolated cell fractions were generated using FlowJo software v10.

#### scRNA-seq library preparation, sequencing, alignment, quality control, and data analysis

5,000 to 10,000 cells isolated by flow cytometry were loaded onto a Chromium Controller (10X Genomics), followed by library preparation according to the manufacturer’s recommendations (Single Cell 3’ V3 assay) and sequencing on a HiSeq4000 instrument (Illumina). Libraries were de-multiplexed, aligned to the mm10 mouse transcriptome, and unique molecular identifiers (UMIs) were counted using Cell Ranger (10X Genomics) v3.1.1. Data were analyzed using the Seurat package v4.0.3 in R (Butler et al., 2018). Cells with ≥1,000 and ≤ 4,000 detected genes, >2,500 and <15,000 total transcripts, and <10% mitochondrial transcripts were retained. Merged datasets utilized the “merge” function and data were normalized and log-transformed using the “SCTransform” function in Seurat, regressing out mitochondrial read fractions and other confounding variables. Datasets integrated after normalization used the “integrate” function in Seurat. Differential markers were identified in clusters using the “FindAllMarkers” function in Seurat, with parameters: min.pct 0.25 and logfc.threshold 0.25. To identify cell types in each collection, the data were queried for known mesenchymal cell-specific genes(Kinchen et al., 2018; McCarthy et al., 2020b). UMAP plots for gene expression were generated using the “FeaturePlot” function.

For merged datasets (Figures 3A and S2A), the top 15 principal components were selected using the “FindNeighbors” function, followed by identifying clusters using the “FindClusters” (resolution: 2) function in Seurat. The “RunUMAP” function was used to reduce the top 15 principal components using the “uwot” method. Based on marker expression, clusters were merged to compile 20 mesenchymal cell populations; among these, we removed *Mki67^+^* (proliferating), *Ptprc^+^*(leukocytes), and *Ccl19^+^* (follicle reticular, FRC) cells (Figure S2A) to generate the final merged dataset (Figure 3A). Separately, *Ptprc^+^*and *Mki67^+^* cell subsets were annotated on the basis of published markers (Xu et al., 2019) and those listed in Figure S2A, respectively. To generate correlation heatmaps (Figure 3C) and gene lists (Table S2), we separated SM cell types using the “FindAllMarkers” function in Seurat and plotted the top 10 and top 50 marker genes for each. Genes differentially expressed between P14 and P1/P2 (Figure 4B) were identified for each cell type using the “FindMarkers” function and dot plots depicting log_2_ fold-differences were generated using code based on the “DotPlot” function within the ggplot2 package in R. Heatmaps show average cell expression of the top 50 differentially expressed genes at each harvest between P1 or P2 and P14 (Figure S4B). Integrated analysis (Figure S5F) included published adult scRNA-seq data(McCarthy et al., 2020b) (Gene Expression Omnibus, GSE130681).

#### *In vitro* co-cultures, imaging, quantitation, and analysis

Unfractionated mesenchyme extracted from P14 or adult *Pdgfra^H2B-eGFP^*mice was plated on non-pyrogenic, gas plasma surface-treated polystyrene tissue culture plates (Falcon) in Dulbecco’s Modified Eagle and F12 media (Gibco, 12634-010) supplemented with penicillin, streptomycin, Glutamax, HEPES buffer, and 10% FBS (Basal media + FBS, see Ref 35 (McCarthy et al., 2020b) for extraction methods, Figure S1C). Using forceps, we carefully peeled off SI muscularis propria (MP), and in the colon, the muscularis mucosae from *Pdgfra^H2B-eGFP^*mice. SM was disaggregated for 10 min in HBSS containing 5%FBS and 3 mg/mL collagenase II, then plated as described above. The medium was replaced 24 h after plating; 2 to 3 days later, cells were removed using 0.25% Trypsin/EDTA (Corning), washed in FACS buffer, and GFP^hi^ (ISEMFs), CD81^+^Pdgfra^lo^ (trophocytes), CD81^-^Pdgfra^lo^ stroma, and GFP^-^ cell fractions were harvested by flow cytometry (Figure S1C) as described previously(McCarthy et al., 2020b) as unfractionated mesenchyme (Figure 2A) or SM populations (Figure 7C). MP was disaggregated using 0.25% trypsin/EDTA for 5 min, washed with PBS/0.1%BSA, and placed directly into co-cultures with isolated crypts.

For organoid co-cultures (Figures 2A and S1E), crypts were plated for 2 days in complete basal media supplemented with N2, B27 and N-acetylcysteine as described(Sato and Clevers, 2013), EGF (Thermo Fisher, 50 ng/mL), RSPO1 (10% culture supernatant from 293T-HA-RspoI-Fc cells), and rNOG (Peprotech, 100 ng/mL), then removed from Matrigel in Cell Recovery Solution (Corning) for 10 min at 4°C, and recovered by centrifugation at 200 *g* for 15 min. 50-100 crypts were replated in 20 μl Matrigel drops in 24-well tissue culture plates, together with 2×10^4^ mesenchymal cells harvested, cultured, and purified as described above. Complete Basal Media was replaced containing EGF, RSPO1 (10% culture supernatant from 293T-HA-RspoI-Fc cells), and rNOG, rGREM1 (Thermo Fisher, 100 ng/ml), or BMP2 (Peprotech #120-02, 50 ng/mL) and BMP7 (R&D#5666-BP, 50 ng/mL). Bright field and fluorescent organoid images were captured using an Olympus CKX53 or Nikon Eclipse T/2 microscope, respectively. Organoid features were scored from 3 or 4 experiments with 2 or 3 technical replicates per group per experiment. RNA was harvested at 48 h and qRT-PCR was performed on 3 or 4 replicates from each treatment group; mRNA expression values are represented relative to Gapdh (2^-ΔCT^) and relative to organoids cultured without additional cells.

For culture experiments with BMPi (Figure 7A), ∼100 isolated crypts were plated in Matrigel drops in media containing a sub-optimal concentration of RSPO1 (0.5% culture supernatant from 293T-HA-Rspo1-Fc cells), EGF (50 ng/mL), and indicated concentrations of rNOG, rGREM1, or rGREM2 (R&D#2069-PR). For culture experiments with RSPOs (Figures 7E and S7I), ∼100 isolated crypts were plated in Matrigel drops in basal condition media containing rNOG (100 ng/mL), EGF (50 ng/mL), and the indicated concentrations of mRSPO2, mRSPO3 (R&D, 6946-RS and 4120-RS), hRSPO1, hRSPO2, or hRSPO3 (R&D, 4645-RS, 3266-RS, and 3500-RS). Organoid structures were counted with a Nikon Eclipse TS100 microscope 5 days after plating. Three biological replicates are reported per condition, each including 3 technical replicates. Figure S6C and 7G were compiled as a composite using Microsoft 365 ProPlus Office PowerPoint.

#### Statistics and reproducibility

Statistical analyses were performed in Prism software package v7.03 (GraphPad). Replicate numbers, statistical methods, and P values are given in the respective figure legends. No sample size estimations and no blinding were performed.

#### Data and code availability

Data are deposited in the Gene Expression Omnibus (https://www.ncbi.nlm.nih.gov/geo/query/acc.cgi?acc=GSE184158). Reviewers may access data using the token **apqtymkwpnwnlmx**. No new software was developed for this study.

## SUPPLEMENTAL FIGURE LEGENDS

**Figure S1. pSMAD1/5 and ISEMF distributions during pre- and post-natal development, organoid co-culture features, and isolation of mesenchymal populations;** related to Figures 1 and 2.

**A)** Epithelial nuclear pSMAD1/5 (arrowheads) increases over time at villus bottoms. Magenta: pSMAD1/5, green: Laminin (basement membrane), blue: DAPI, cr: crypt, scale bars: 100 μm.

**B)** Representative immunostaining with the indicated antibodies in *Pdgfra^H2B-eGFP^* fetuses at embryonic days (E) 14.5 and E16.5. GFP^hi^ ISEMFs initially concentrate at emerging villus tips, while GFP^lo^ cells are present throughout the stroma. Top and bottom rows show the same fields with (top) or without (bottom) DAPI signals. n = 3 embryos per stage, scale bar 50 μm, dashed white lines: inter-villus epithelium.

**C)** Schema for mesenchyme isolation and co-culture. P14 or adult (>8 weeks) *Pdgfra^H2B-eGFP^* whole mesenchyme was plated, sorted 2-3 days later by flow cytometry (FACS), and different fractions were placed in co-culture with isolated crypt epithelium. FACS plots show GFP^hi^, GFP^lo^, and GFP^-^ fractions; histograms show GFP^lo^ cell staining with CD81 antibody.

**D)** Organoids imaged on consecutive days after ISEMF co-culture show retraction of budding structures (arrowheads).

**E)** Two-day old organoids cultured with recombinant RSPO1 (R), EGF (E), and either NOG (N) or GFP^hi^ ISEMFs from P14 *Pdgfra^H2B-eGFP^* mouse SI. Representative organoids are shown 48 h later (scale bar 100 μm). NOG rescues failure to bud in crypts co-cultured with P14 ISEMFs (arrowheads). Graph displays data from 4 biological replicates. Statistical differences, determined by one-way ANOVA followed by Tukey’s multiple comparisons test, are reported relative to +ER (no cells). ns: not significant, **p<0.01.

**Figure S2. Merged postnatal scRNA datasets from SI mesenchyme;** related to Figure 3.

**A)** Clustering of cell populations by uniform manifold approximation and projection (UMAP, left) and markers signifying distinct cell types (right) in merged postnatal scRNA-seq datasets.

**B)** *Ptprc^+^* cells isolated from the combined dataset (UMAP, left) annotated for distinct immune cells and leukocytes based on marker expression (right).

**C)** Relative fractions of proliferative (*Mki67^+^*) cell clusters at different postnatal ages.

**D)** Cell distributions within the global UMAP plot with respect to postnatal age (left) or method of mesenchymal cell isolation (right, unfractionated vs. GFP^+^ cells).

**E)** Cell types at each postnatal age. Compared to other populations, MM and LPM cells (bold) increase substantially between P1 and P14.

**Figure S3: ISH images and quantitation of SM cell-type marker expression;** related to Figure 3.

**A)** Negative control ISH probe (RNAscope universal control: *B. subtilis DapB* gene) hybridized to P14 SI. Left, color image; right, greyscale version of the same image. All scale bars 50 μm.

**B)** Fractions of SI sub-epithelial SM cell types showing *Nog*, *Pcp4* or *Osr1* expression at P14. Parentheses indicate cell numbers counted (denominators for the fractions) in SI from 2 mice (“submucosa” refers to the trophocyte zone between MM and MP; “plexus” refers to intermuscular neurons).

**C)** *Hhip* expression across P14 SM populations determined by scRNA-seq (violin plot) and ISH (RNAscope images). *Hhip*, absent in MP layers, is present in *Grem2^-^* LPM (dashed red box) and *Grem2^+^*MM (dashed white box – MM and LPM are oriented perpendicular to each other), both magnified on the right. Greyscale versions of the same images are shown to the right of (LPM) or below (MM) color ISH images. Arrowheads: *Hhip* and *Grem2* co-expressing MM.

**D)** *Grem1* ISH at P14 reveals expression in MM and/or select MP layers, as quantified to the right in SI from 2 mice (parentheses in the table indicate cell numbers counted). Green: LAMININ, magenta: *Grem1* (greyscale version of the same image is shown below), white/green dashed lines: MP, arrowheads: MM, cr: crypts.

**E)** Double ISH for *Nog* and *Grem2* highlights that *Grem2^+^* MM is distinct from *Nog^+^* superficial MP. Grey: LAMININ, magenta: *Nog*, green: *Grem2* (greyscale versions without LAM signal shown to the right), arrowheads: expressing (filled) and non-expressing (blank) cells.

**Figure S4. Differential postnatal gene expression and CellPhone DB analysis;** related to Figures 3 and 4.

**A)** Average expression of Wnt, secreted Wnt antagonist, and BMP genes in combined postnatal intestinal mesenchyme dataset (Figure 3A).

**B)** Average expression of the top 50 genes differentially expressed between P1/2 (aggregated) and P14, shown at those ages, P4+P5 (aggregated), and P9 in trophocytes (left) and *Nog^+^* superficial MP (right).

**Figure S5. Analysis of postnatal *Pdgfra^H2B-eGFP^* and *Pdgfra^CreERT2^;Rosa26^tdTom^* and adult MM populations;** related to Figures 4 and 5.

**A)** Fractions of GFP^+^ MM or LPM cells in *Pdgfra^H2B-eGFP^* intestines at the indicated ages. GFP^+^ cells are in a majority until at least P14, but both adult populations are GFP^-^. 3 independent animals per timepoint, n: total cells counted in each replicate (rep).

**B)** Markers and graph-based UMAP clustering in GFP^+^ cell fractions from P14 (top) and average BMPi and Rspo gene expression at P14 (bottom).

**C)** *Pdgfra^Cre(ER-T2)^;Rosa26^TdTom^*pups were treated with tamoxifen (TAM) at P0 and intestines were examined at P1. In the representative image, both PDGFRA^+^ ISEMFs (PDGFRA antibody labels ISEMFs green, arrowhead) and undefined stroma (arrow, red PDGFRA^lo^ cells above ACTA2+ yellow cells) express TdTom after TAM-mediated Cre activation. Scale bar, 50 μm.

**D)** Delineation of mesenchymal cell populations in adult *Pdgfra^H2B-eGFP^* duodenum. ACTA2 (smooth muscle) and CD31 (blood vessels) Ab stains identify thin MM (arrowheads) and LPM, distinct from the thick MP. Dashed white line: superficial MP, cr: crypts.

**E)** scRNA analysis of 2,224 adult *Myh11^+^*SI cells and graph-based UMAP clustering of cell populations. *Hhip* (red) expression principally marks LPM; *Grem2* (blue) marks the substantially fewer recoverable MM cells.

**F)** scRNA data from *Myh11^+^*SI cells (**e**) were combined with scRNA data from unfractionated adult SI mesenchyme^(McCarthy^ ^et^ ^al., 2020b)^ (GEO series GSE130681). Molecular markers identified each cell type (see Figure S2A). Average BMPi and Rspo gene expression and cell numbers in each cluster are shown to the right.

**Figure S6. ISH in Adult SI and SM ablation in Myh11;DTA mice;** related to Figure 5.

**A-B)** Representative ISH of adult *Pdgfra^H2B-eGFP^* SI with probes for *Rspo3* (**A**, magenta and below greyscale), *Grem1* and *Grem2* (**B**, red and yellow respectively). In line with scRNA data (Figure 4A), *Rspo3* is expressed in MM (arrowheads), MP (dotted white lines demarcate superficial MP), and scattered submucosal cells. In addition to these cells, *Grem1* is expressed in GFP^+^ sub-cryptal trophocytes (green arrowheads in **B**). *Grem2* is expressed in MM, superficial MP, and interstitial cells of Cajal (ICC; cr: crypts).

**C)** Composite *Grem1* ISH overlayed onto IHC-stained *Pdgfra^H2B-eGFP^* adult mouse duodenum (sections 10 μm removed from each other), showing high expression in both MM (arrowhead) and superficial MP (between brackets; PDGFRA in red, *Pdgfra^H2B-eGFP^* in green, DAPI in blue, LAMININ in yellow).

**D)** Representative image of a *Myh11^Cre(ER-T2)^; Pdgfra^H2B-eGFP^*;*R26R^TdTom^* mouse injected at P14, harvested at P21, and stained for ACTA2 (blue). Cre activity (TdTom, magenta) and ACTA2 expression coincide (n= 3 mice). All scale bars: 50 μm.

**Figure S7. Investigation of SM functions and potency of human RSPO;** related to Figures 6 and 7.

**A)** Reduced body weight and SI length at P16 and P21 after TAM treatment of *Myh11^Cre(ER-T2)^;R26R^DTA^* pups at P14. Images show examples at P21. n= 7 or 8 animals, ***p <0.001, ****p <0.0001 using unpaired Student’s t-test.

**B)** Quantitation of circular and longitudinal MP cells at P16 and P21 in control and *Myh11^Cre(ER-T2)^;R26R^DTA^* mice after TAM treatment at P14. n=3 or 4 animals, >4 mm examined per SI. *p <0.05 using unpaired Student’s t-test.

**C)** Crosses with *Pdgfra^H2B-eGFP^* mice show a small decrease in sub-cryptal *Pdgfra^lo^* cells (arrowheads), likely reflecting loss of *Myh11^+^* MM and LPM precursors. n=3 animals, >4 mm examined in each, *p <0.05 using unpaired Student’s t-test.

**D)** Crypt cell proliferation, assessed by BrdU injection 1 h before euthanasia, is similar in TAM-injected control and *Myh11;DTA* mice. n=3 animals in each cohort, ≥25 SI crypts per animal, differences are non-significant by unpaired Student’s t-test. Scale bar 50 μm.

**E)** F-actin (phalloidin)-stained SI epithelium whole-mounts show fewer bifid crypts at P16 in *Myh11^Cre(ER-T2)^;Rosa26^DTA^* pups injected with TAM at P14. Scale bar 50 μm.

**F)** Neither P14 nor adult MP co-cultures with P14 SI crypts elicit organoid growth but both yield robust organoids in the presence of sub-optimal rEGF (E) and rRSPO (R) concentrations. Organoid numbers are represented relative 76.7 ±12.9 organoids in ENR medium (n=3). ****p <0.0001 using Student’s t-test.

**G)** SI crypts cultured over manually isolated colonic MM (dashed white line) develop within 1 day into spheroidal structures (n=3). The image is a collage assembled from multiple microscopic fields (scale bar 200 μm). Inset magnifies one such field (scale bar 100 μm).

**H)** Representative FACS histogram of GFP^+^ cells isolated from manually stripped colonic MM and examined for CD81 expression. -Ab: control flow cytometry without CD81 Ab.

**I)** Adult SI crypts cultured in media containing murine rEGF, rNOG, and different concentrations of the indicated human RSPO (n=3 biological replicates each). Organoid numbers are graphed with respect to 146.3 ±20.6 organoids formed in 100 ng/ml hRSPO3. Statistical significance was determined using one-way ANOVA followed by Tukey’s posttest. ***P< 0.001, *P <0.05.

## REFERENCES

Al-Nafussi, A.I., and Wright, N.A. (1982). Cell kinetics in the mouse small intestine during immediate postnatal life. Virchows Arch B Cell Pathol Incl Mol Pathol 40, 51–62.

Auclair, B.A., Benoit, Y.D., Rivard, N., Mishina, Y., and Perreault, N. (2007). Bone morphogenetic protein signaling is essential for terminal differentiation of the intestinal secretory cell lineage. Gastroenterology 133, 887–896.

Bahar Halpern, K., Massalha, H., Zwick, R.K., Moor, A.E., Castillo-Azofeifa, D., Rozenberg, M., Farack, L., Egozi, A., Miller, D.R., Averbukh, I., et al. (2020). Lgr5+ telocytes are a signaling source at the intestinal villus tip. Nat Commun 11, 1936.

Barker, N., van Es, J.H., Kuipers, J., Kujala, P., van den Born, M., Cozijnsen, M., Haegebarth, A., Korving, J., Begthel, H., Peters, P.J., et al. (2007). Identification of stem cells in small intestine and colon by marker gene Lgr5. Nature 449, 1003–1007.

Batts, L.E., Polk, D.B., Dubois, R.N., and Kulessa, H. (2006). Bmp signaling is required for intestinal growth and morphogenesis. Developmental dynamics : an official publication of the American Association of Anatomists 235, 1563–1570.

Bernier-Latmani, J., and Petrova, T.V. (2016). High-resolution 3D analysis of mouse small-intestinal stroma. Nat Protoc 11, 1617–1629.

Brugger, M.D., Valenta, T., Fazilaty, H., Hausmann, G., and Basler, K. (2020). Distinct populations of crypt-associated fibroblasts act as signaling hubs to control colon homeostasis. PLoS Biol 18, e3001032.

Bry, L., Falk, P., Huttner, K., Ouellette, A., Midtvedt, T., and Gordon, J.I. (1994). Paneth cell differentiation in the developing intestine of normal and transgenic mice. Proceedings of the National Academy of Sciences of the United States of America 91, 10335–10339.

Butler, A., Hoffman, P., Smibert, P., Papalexi, E., and Satija, R. (2018). Integrating single-cell transcriptomic data across different conditions, technologies, and species. Nature biotechnology 36, 411–420.

Calvert, R., and Pothier, P. (1990). Migration of fetal intestinal intervillous cells in neonatal mice. Anat Rec 227, 199–206.

Chee, Y.C., Pahnke, J., Bunte, R., Adsool, V.A., Madan, B., and Virshup, D.M. (2018). Intrinsic Xenobiotic Resistance of the Intestinal Stem Cell Niche. Developmental cell 46, 681–695 e685.

Chen, L., Toke, N.H., Luo, S., Vasoya, R.P., Fullem, R.L., Parthasarathy, A., Perekatt, A.O., and Verzi, M.P. (2019). A reinforcing HNF4-SMAD4 feed-forward module stabilizes enterocyte identity. Nat Genet 51, 777–785.

Chen, P.H., Chen, X., Lin, Z., Fang, D., and He, X. (2013). The structural basis of R-spondin recognition by LGR5 and RNF43. Genes & development 27, 1345–1350.

Cheng, H., and Bjerknes, M. (1985). Whole population cell kinetics and postnatal development of the mouse intestinal epithelium. Anat Rec 211, 420–426.

Chi, P., Chen, Y., Zhang, L., Guo, X., Wongvipat, J., Shamu, T., Fletcher, J.A., Dewell, S., Maki, R.G., Zheng, D., et al. (2010). ETV1 is a lineage survival factor that cooperates with KIT in gastrointestinal stromal tumours. Nature 467, 849–853.

Chung, M.I., Bujnis, M., Barkauskas, C.E., Kobayashi, Y., and Hogan, B.L.M. (2018). Niche-mediated BMP/SMAD signaling regulates lung alveolar stem cell proliferation and differentiation. Development 145.

Clevers, H. (2013). The intestinal crypt, a prototype stem cell compartment. Cell 154, 274–284.

Cretoiu, D., Cretoiu, S.M., Simionescu, A.A., and Popescu, L.M. (2012). Telocytes, a distinct type of cell among the stromal cells present in the lamina propria of jejunum. Histol Histopathol 27, 1067–1078.

Davis, H., Irshad, S., Bansal, M., Rafferty, H., Boitsova, T., Bardella, C., Jaeger, E., Lewis, A., Freeman-Mills, L., Giner, F.C., et al. (2015). Aberrant epithelial GREM1 expression initiates colonic tumorigenesis from cells outside the stem cell niche. Nat Med 21, 62–70.

Eilken, H.M., Dieguez-Hurtado, R., Schmidt, I., Nakayama, M., Jeong, H.W., Arf, H., Adams, S., Ferrara, N., and Adams, R.H. (2017). Pericytes regulate VEGF-induced endothelial sprouting through VEGFR1. Nat Commun 8, 1574.

Farin, H.F., Van Es, J.H., and Clevers, H. (2012). Redundant sources of Wnt regulate intestinal stem cells and promote formation of Paneth cells. Gastroenterology 143, 1518–1529 e1517.

Fordham, R.P., Yui, S., Hannan, N.R., Soendergaard, C., Madgwick, A., Schweiger, P.J., Nielsen, O.H., Vallier, L., Pedersen, R.A., Nakamura, T., et al. (2013). Transplantation of expanded fetal intestinal progenitors contributes to colon regeneration after injury. Cell stem cell 13, 734–744.

Greicius, G., Kabiri, Z., Sigmundsson, K., Liang, C., Bunte, R., Singh, M.K., and Virshup, D.M. (2018). PDGFRalpha(+) pericryptal stromal cells are the critical source of Wnts and RSPO3 for murine intestinal stem cells in vivo. Proceedings of the National Academy of Sciences of the United States of America 115, E3173–E3181.

Hamilton, T.G., Klinghoffer, R.A., Corrin, P.D., and Soriano, P. (2003). Evolutionary divergence of platelet-derived growth factor alpha receptor signaling mechanisms. Molecular and cellular biology 23, 4013–4025.

Haramis, A.P., Begthel, H., van den Born, M., van Es, J., Jonkheer, S., Offerhaus, G.J., and Clevers, H. (2004). De novo crypt formation and juvenile polyposis on BMP inhibition in mouse intestine. Science 303, 1684–1686.

He, X.C., Zhang, J., Tong, W.G., Tawfik, O., Ross, J., Scoville, D.H., Tian, Q., Zeng, X., He, X., Wiedemann, L.M., et al. (2004). BMP signaling inhibits intestinal stem cell self-renewal through suppression of Wnt-beta-catenin signaling. Nature genetics 36, 1117–1121.

Howe, J.R., Bair, J.L., Sayed, M.G., Anderson, M.E., Mitros, F.A., Petersen, G.M., Velculescu, V.E., Traverso, G., and Vogelstein, B. (2001). Germline mutations of the gene encoding bone morphogenetic protein receptor 1A in juvenile polyposis. Nature genetics 28, 184–187.

Howe, J.R., Roth, S., Ringold, J.C., Summers, R.W., Jarvinen, H.J., Sistonen, P., Tomlinson, I.P., Houlston, R.S., Bevan, S., Mitros, F.A., et al. (1998). Mutations in the SMAD4/DPC4 gene in juvenile polyposis. Science 280, 1086–1088.

Huycke, T.R., Miller, B.M., Gill, H.K., Nerurkar, N.L., Sprinzak, D., Mahadevan, L., and Tabin, C.J. (2019). Genetic and Mechanical Regulation of Intestinal Smooth Muscle Development. Cell 179, 90–105 e121.

Itzkovitz, S., Blat, I.C., Jacks, T., Clevers, H., and van Oudenaarden, A. (2012). Optimality in the development of intestinal crypts. Cell 148, 608–619.

Jaeger, E., Leedham, S., Lewis, A., Segditsas, S., Becker, M., Cuadrado, P.R., Davis, H., Kaur, K., Heinimann, K., Howarth, K., et al. (2012). Hereditary mixed polyposis syndrome is caused by a 40-kb upstream duplication that leads to increased and ectopic expression of the BMP antagonist GREM1. Nature genetics 44, 699–703.

Kanda, T., Sullivan, K.F., and Wahl, G.M. (1998). Histone-GFP fusion protein enables sensitive analysis of chromosome dynamics in living mammalian cells. Curr Biol 8, 377–385.

Karlsson, L., Lindahl, P., Heath, J.K., and Betsholtz, C. (2000). Abnormal gastrointestinal development in PDGF-A and PDGFR-(alpha) deficient mice implicates a novel mesenchymal structure with putative instructive properties in villus morphogenesis. Development 127, 3457–3466.

Kim, J.E., Fei, L., Yin, W.C., Coquenlorge, S., Rao-Bhatia, A., Zhang, X., Shi, S.S.W., Lee, J.H., Hahn, N.A., Rizvi, W., et al. (2020). Single cell and genetic analyses reveal conserved populations and signaling mechanisms of gastrointestinal stromal niches. Nat Commun 11, 334.

Kim, T.H., Escudero, S., and Shivdasani, R.A. (2012). Intact function of Lgr5 receptor-expressing intestinal stem cells in the absence of Paneth cells. Proceedings of the National Academy of Sciences of the United States of America 109, 3932–3937.

Kinchen, J., Chen, H.H., Parikh, K., Antanaviciute, A., Jagielowicz, M., Fawkner-Corbett, D., Ashley, N., Cubitt, L., Mellado-Gomez, E., Attar, M., et al. (2018). Structural remodeling of the human colonic mesenchyme in inflammatory bowel disease. Cell 175, 372–386 e317.

Koppens, M.A.J., Davis, H., Valbuena, G.N., Mulholland, E.J., Nasreddin, N., Colombe, M., Antanaviciute, A., Biswas, S., Friedrich, M., Lee, L., et al. (2021). Bone Morphogenetic Protein Pathway Antagonism by Grem1 Regulates Epithelial Cell Fate in Intestinal Regeneration. Gastroenterology 161, 239–254 e239.

Kosinski, C., Li, V.S., Chan, A.S., Zhang, J., Ho, C., Tsui, W.Y., Chan, T.L., Mifflin, R.C., Powell, D.W., Yuen, S.T., et al. (2007). Gene expression patterns of human colon tops and basal crypts and BMP antagonists as intestinal stem cell niche factors. Proceedings of the National Academy of Sciences of the United States of America 104, 15418–15423.

Kurahashi, M., Nakano, Y., Hennig, G.W., Ward, S.M., and Sanders, K.M. (2012a). Platelet-derived growth factor receptor alpha-positive cells in the tunica muscularis of human colon. Journal of cellular and molecular medicine 16, 1397–1404.

Kurahashi, M., Nakano, Y., Hennig, G.W., Ward, S.M., and Sanders, K.M. (2012b). Platelet-derived growth factor receptor alpha-positive cells in the tunica muscularis of human colon. J Cell Mol Med 16, 1397–1404.

Langlands, A.J., Almet, A.A., Appleton, P.L., Newton, I.P., Osborne, J.M., and Nathke, I.S. (2016). Paneth Cell-Rich Regions Separated by a Cluster of Lgr5+ Cells Initiate Crypt Fission in the Intestinal Stem Cell Niche. PLoS Biol 14, e1002491.

Link, A., Vogt, T.K., Favre, S., Britschgi, M.R., Acha-Orbea, H., Hinz, B., Cyster, J.G., and Luther, S.A. (2007). Fibroblastic reticular cells in lymph nodes regulate the homeostasis of naive T cells. Nat Immunol 8, 1255–1265.

Madisen, L., Zwingman, T.A., Sunkin, S.M., Oh, S.W., Zariwala, H.A., Gu, H., Ng, L.L., Palmiter, R.D., Hawrylycz, M.J., Jones, A.R., et al. (2010). A robust and high-throughput Cre reporting and characterization system for the whole mouse brain. Nat Neurosci 13, 133–140.

Martin-Alonso, M., Iqbal, S., Vornewald, P.M., Lindholm, H.T., Damen, M.J., Martinez, F., Hoel, S., Diez-Sanchez, A., Altelaar, M., Katajisto, P., et al. (2021). Smooth muscle-specific MMP17 (MT4-MMP) regulates the intestinal stem cell niche and regeneration after damage. Nat Commun 12, 6741.

Maskens, A.P., and Dujardin-Loits, R.M. (1981). Kinetics of tissue proliferation in colorectal mucosa during post-natal growth. Cell Tissue Kinet 14, 467–477.

McCarthy, N., Kraiczy, J., and Shivdasani, R.A. (2020a). Cellular and molecular architecture of the intestinal stem cell niche. Nature cell biology 22, 1033–1041.

McCarthy, N., Manieri, E., Storm, E.E., Saadatpour, A., Luoma, A.M., Kapoor, V.N., Madha, S., Gaynor, L.T., Cox, C., Keerthivasan, S., et al. (2020b). Distinct Mesenchymal Cell Populations Generate the Essential Intestinal BMP Signaling Gradient. Cell stem cell 26, 391–402 e395.

Nerurkar, N.L., Mahadevan, L., and Tabin, C.J. (2017). BMP signaling controls buckling forces to modulate looping morphogenesis of the gut. Proc Natl Acad Sci USA 114, 2277–2282.

Peng, W.C., de Lau, W., Forneris, F., Granneman, J.C., Huch, M., Clevers, H., and Gros, P. (2013). Structure of stem cell growth factor R-spondin 1 in complex with the ectodomain of its receptor LGR5. Cell Rep 3, 1885–1892.

Popescu, L.M., and Faussone-Pellegrini, M.S. (2010). TELOCYTES - a case of serendipity: the winding way from Interstitial Cells of Cajal (ICC), via Interstitial Cajal-Like Cells (ICLC) to TELOCYTES. Journal of cellular and molecular medicine 14, 729–740.

Powell, D.W., Adegboyega, P.A., Di Mari, J.F., and Mifflin, R.C. (2005). Epithelial cells and their neighbors I. Role of intestinal myofibroblasts in development, repair, and cancer. American journal of physiology Gastrointestinal and liver physiology 289, G2–7.

Powell, D.W., Pinchuk, I.V., Saada, J.I., Chen, X., and Mifflin, R.C. (2011). Mesenchymal cells of the intestinal lamina propria. Annual review of physiology 73, 213–237.

Qi, Z., Li, Y., Zhao, B., Xu, C., Liu, Y., Li, H., Zhang, B., Wang, X., Yang, X., Xie, W., et al. (2017). BMP restricts stemness of intestinal Lgr5(+) stem cells by directly suppressing their signature genes. Nat Commun 8, 13824.

Ritsma, L., Ellenbroek, S.I.J., Zomer, A., Snippert, H.J., de Sauvage, F.J., Simons, B.D., Clevers, H., and van Rheenen, J. (2014). Intestinal crypt homeostasis revealed at single-stem-cell level by in vivo live imaging. Nature 507, 362–365.

Roulis, M., and Flavell, R.A. (2016). Fibroblasts and myofibroblasts of the intestinal lamina propria in physiology and disease. Differentiation 92, 116–131.

Sato, T., and Clevers, H. (2013). Primary mouse small intestinal epithelial cell cultures. Methods in molecular biology 945, 319–328.

Sato, T., Vries, R.G., Snippert, H.J., van de Wetering, M., Barker, N., Stange, D.E., van Es, J.H., Abo, A., Kujala, P., Peters, P.J., et al. (2009). Single Lgr5 stem cells build crypt-villus structures in vitro without a mesenchymal niche. Nature 459, 262–265.

Schindelin, J., Arganda-Carreras, I., Frise, E., Kaynig, V., Longair, M., Pietzsch, T., Preibisch, S., Rueden, C., Saalfeld, S., Schmid, B., et al. (2012). Fiji: an open-source platform for biological-image analysis. Nature methods 9, 676–682.

Schmidt, G.H., Winton, D.J., and Ponder, B.A. (1988). Development of the pattern of cell renewal in the crypt-villus unit of chimaeric mouse small intestine. Development 103, 785–790.

Shoshkes-Carmel, M., Wang, Y.J., Wangensteen, K.J., Toth, B., Kondo, A., Massasa, E.E., Itzkovitz, S., and Kaestner, K.H. (2018). Subepithelial telocytes are an important source of Wnts that supports intestinal crypts. Nature 557, 242–246.

St Clair, W.H., and Osborne, J.W. (1985). Crypt fission and crypt number in the small and large bowel of postnatal rats. Cell Tissue Kinet 18, 255–262.

Storm, E.E., Durinck, S., de Sousa e Melo, F., Tremayne, J., Kljavin, N., Tan, C., Ye, X., Chiu, C., Pham, T., Hongo, J.A., et al. (2016). Targeting PTPRK-RSPO3 colon tumours promotes differentiation and loss of stem-cell function. Nature 529, 97–100.

Stzepourginski, I., Nigro, G., Jacob, J.M., Dulauroy, S., Sansonetti, P.J., Eberl, G., and Peduto, L. (2017). CD34+ mesenchymal cells are a major component of the intestinal stem cells niche at homeostasis and after injury. Proceedings of the National Academy of Sciences of the United States of America 114, E506–E513.

Sumigray, K.D., Terwilliger, M., and Lechler, T. (2018a). Morphogenesis and Compartmentalization of the Intestinal Crypt. Developmental cell 45, 183–197 e185.

Sumigray, K.D., Terwilliger, M., and Lechler, T. (2018b). Morphogenesis and compartmentalization of the intestinal crypt. Dev Cell 45, 183–197 e185.

Walton, K.D., Whidden, M., Kolterud, A., Shoffner, S.K., Czerwinski, M.J., Kushwaha, J., Parmar, N., Chandhrasekhar, D., Freddo, A.M., Schnell, S., et al. (2016). Villification in the mouse: Bmp signals control intestinal villus patterning. Development 143, 427–436.

Wang, D., Huang, B., Zhang, S., Yu, X., Wu, W., and Wang, X. (2013). Structural basis for R-spondin recognition by LGR4/5/6 receptors. Genes & development 27, 1339–1344.

Wang, F., Flanagan, J., Su, N., Wang, L.C., Bui, S., Nielson, A., Wu, X., Vo, H.T., Ma, X.J., and Luo, Y. (2012). RNAscope: a novel in situ RNA analysis platform for formalin-fixed, paraffin-embedded tissues. J Mol Diagn 14, 22–29.

Xu, H., Ding, J., Porter, C.B.M., Wallrapp, A., Tabaka, M., Ma, S., Fu, S., Guo, X., Riesenfeld, S.J., Su, C., et al. (2019). Transcriptional Atlas of Intestinal Immune Cells Reveals that Neuropeptide alpha-CGRP Modulates Group 2 Innate Lymphoid Cell Responses. Immunity 51, 696–708 e699.

Yan, K.S., Janda, C.Y., Chang, J., Zheng, G.X.Y., Larkin, K.A., Luca, V.C., Chia, L.A., Mah, A.T., Han, A., Terry, J.M., et al. (2017). Non-equivalence of Wnt and R-spondin ligands during Lgr5(+) intestinal stem-cell self-renewal. Nature 545, 238–242.

Zebisch, M., Xu, Y., Krastev, C., MacDonald, B.T., Chen, M., Gilbert, R.J., He, X., and Jones, E.Y. (2013). Structural and molecular basis of ZNRF3/RNF43 transmembrane ubiquitin ligase inhibition by the Wnt agonist R-spondin. Nat Commun 4, 2787.

