## Supplementary figures and images for "Delineation and birth of a layered intestinal stem cell niche"

### Figure S1

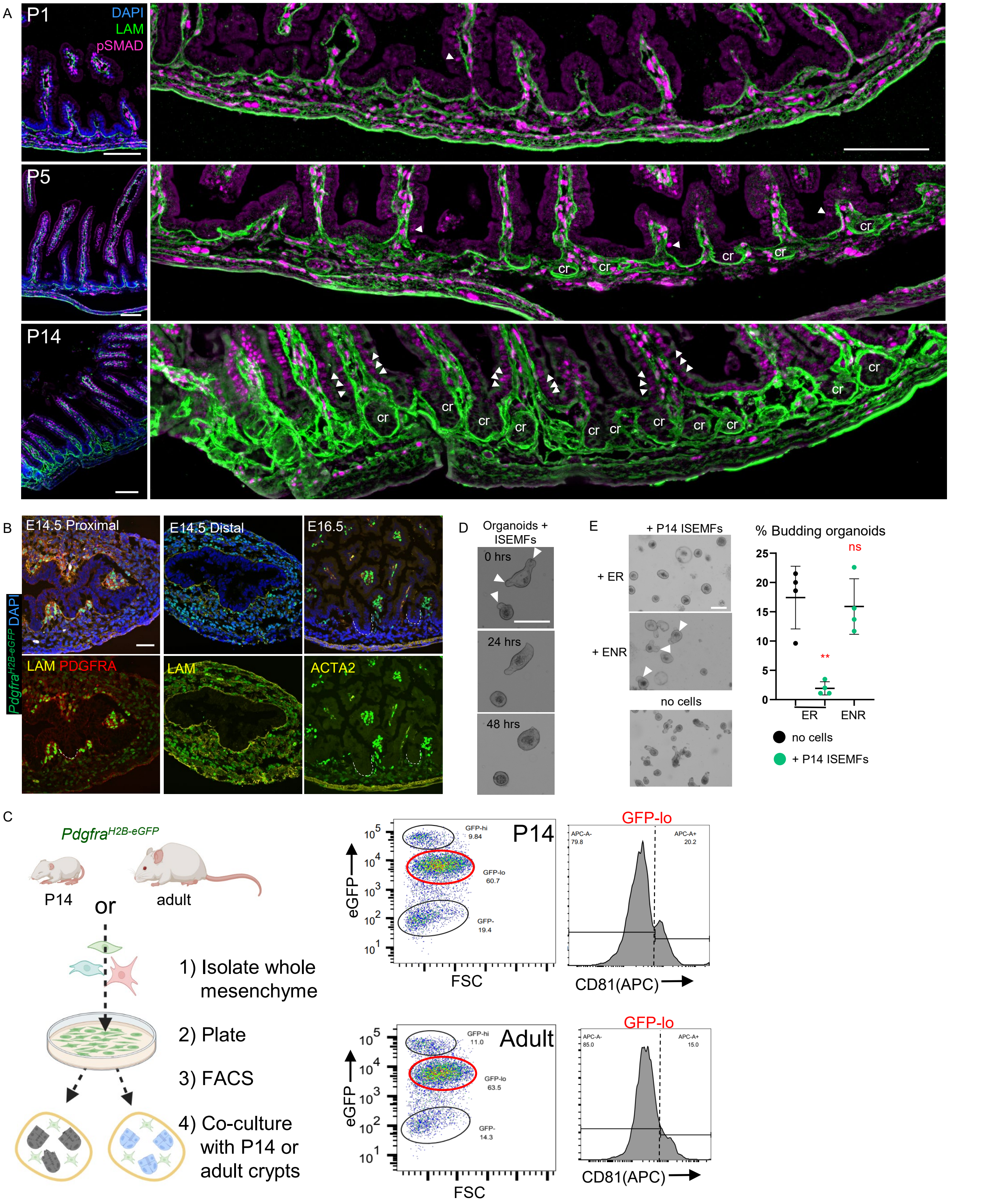

### Figure S4

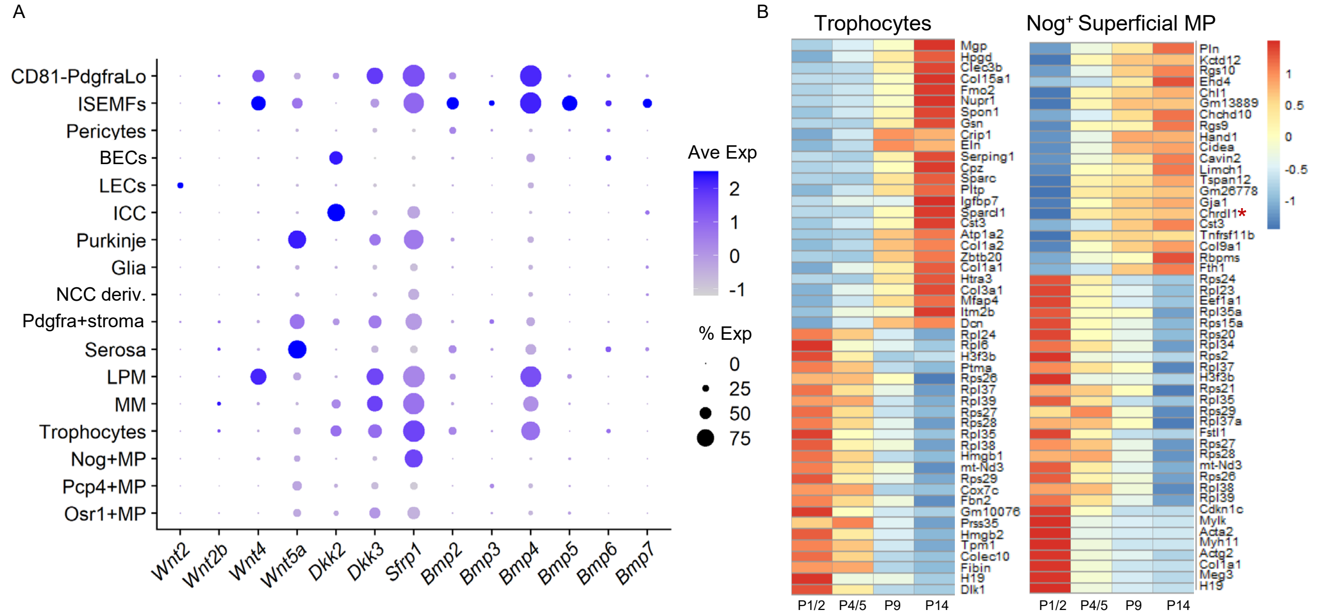

### Figure S5

#### Figure S5

A

GFP⁺ (*Pdgfra*[H2BeGFP]) ACTA2⁺ cells

| | rep | MM % | MM n | LPM % | LPM n |
|---|---|---|---|---|---|
| P1 | 1 | **86** | 35 | **98.1** | 52 |
| | 2 | **100** | 9 | **87.5** | 32 |
| | 3 | **100** | 4 | **96.4** | 28 |
| P5 | 1 | **80** | 54 | **96** | 101 |
| | 2 | **100** | 47 | **75.3** | 69 |
| | 3 | **100** | 82 | **82.5** | 63 |
| P10 | 1 | **94** | 130 | **62.3** | 69 |
| | 2 | **97** | 95 | **65.8** | 85 |
| | 3 | **92** | 133 | **56.9** | 65 |
| P14 | 1 | **87** | 89 | **80.9** | 69 |
| | 2 | **86** | 118 | **69.6** | 85 |
| | 3 | **71** | 78 | **82.2** | 65 |
| Adult | 1 | 0 | 45 | 5.1 | 39 |
| | 2 | 2.3 | 42 | 6.3 | 47 |
| | 3 | 0 | 34 | 0 | 81 |

B

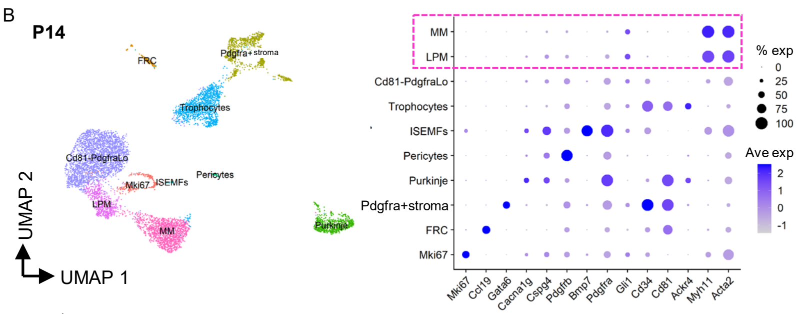


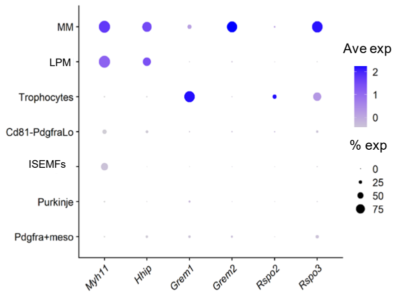


C

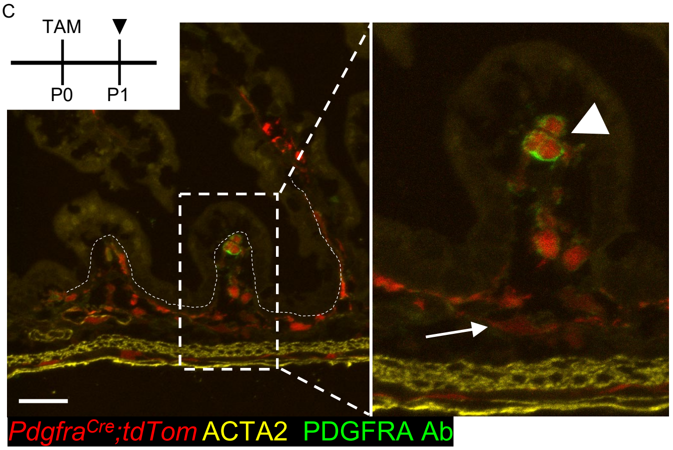


D

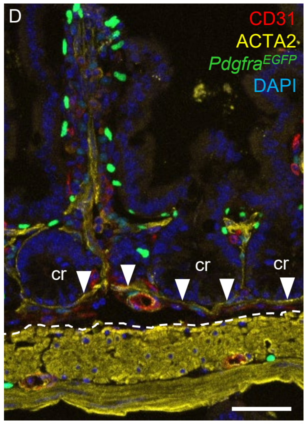


E

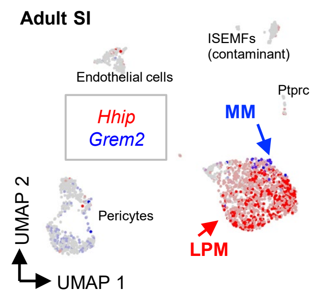


F

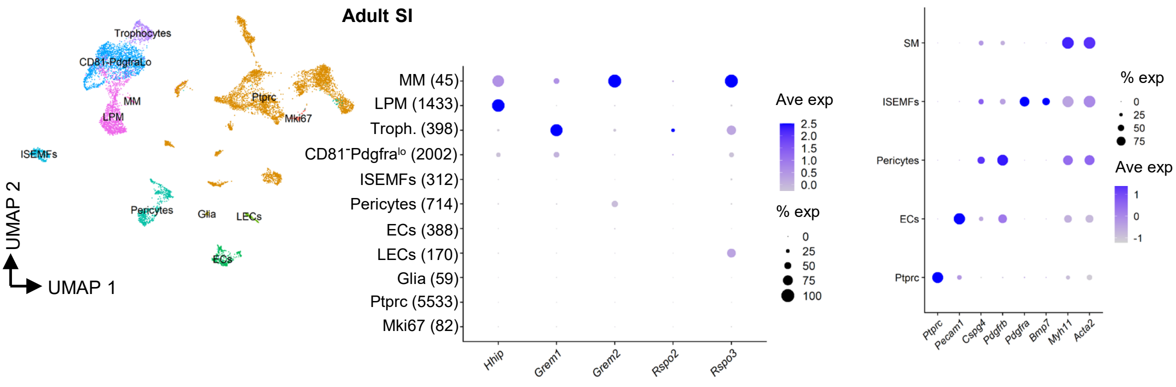

### Figure S6

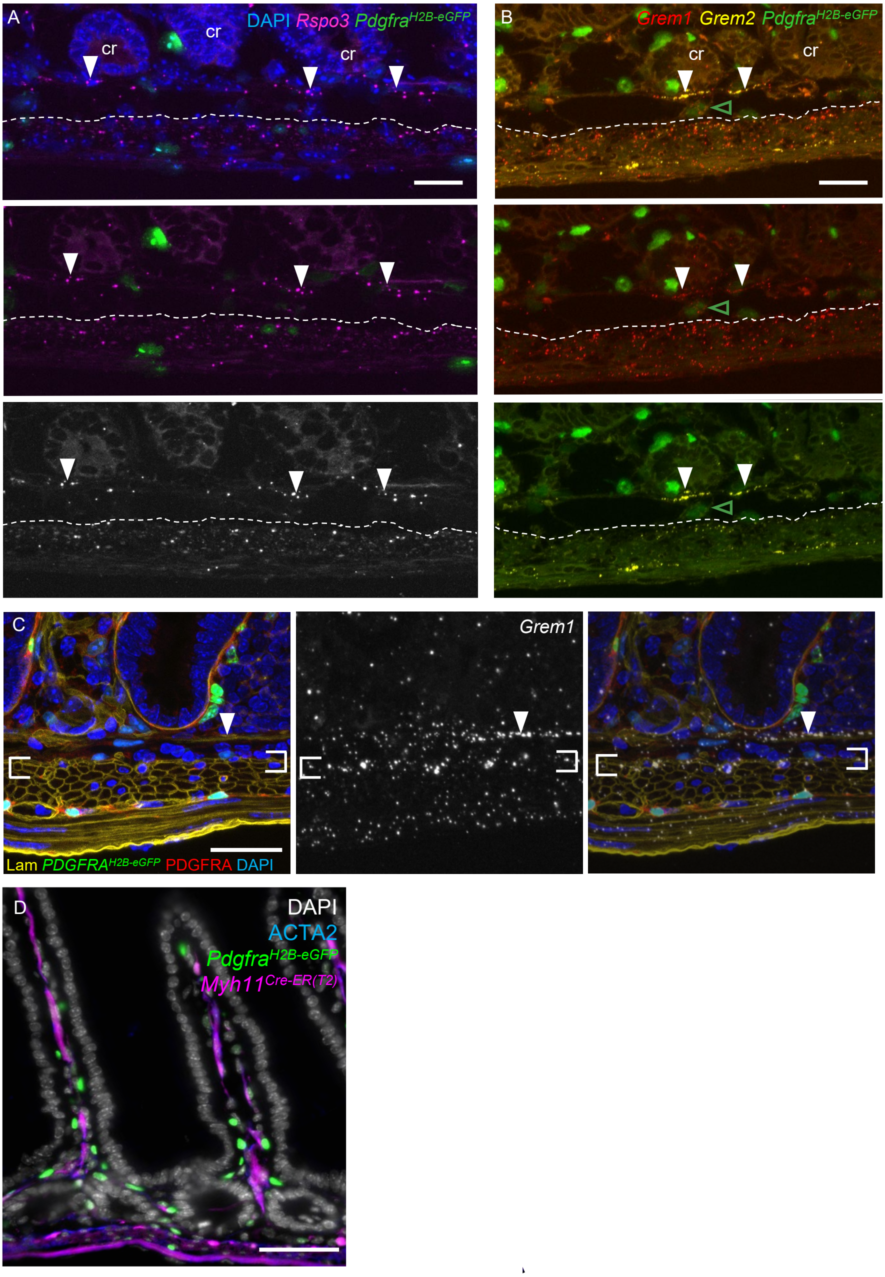

### Figure S7

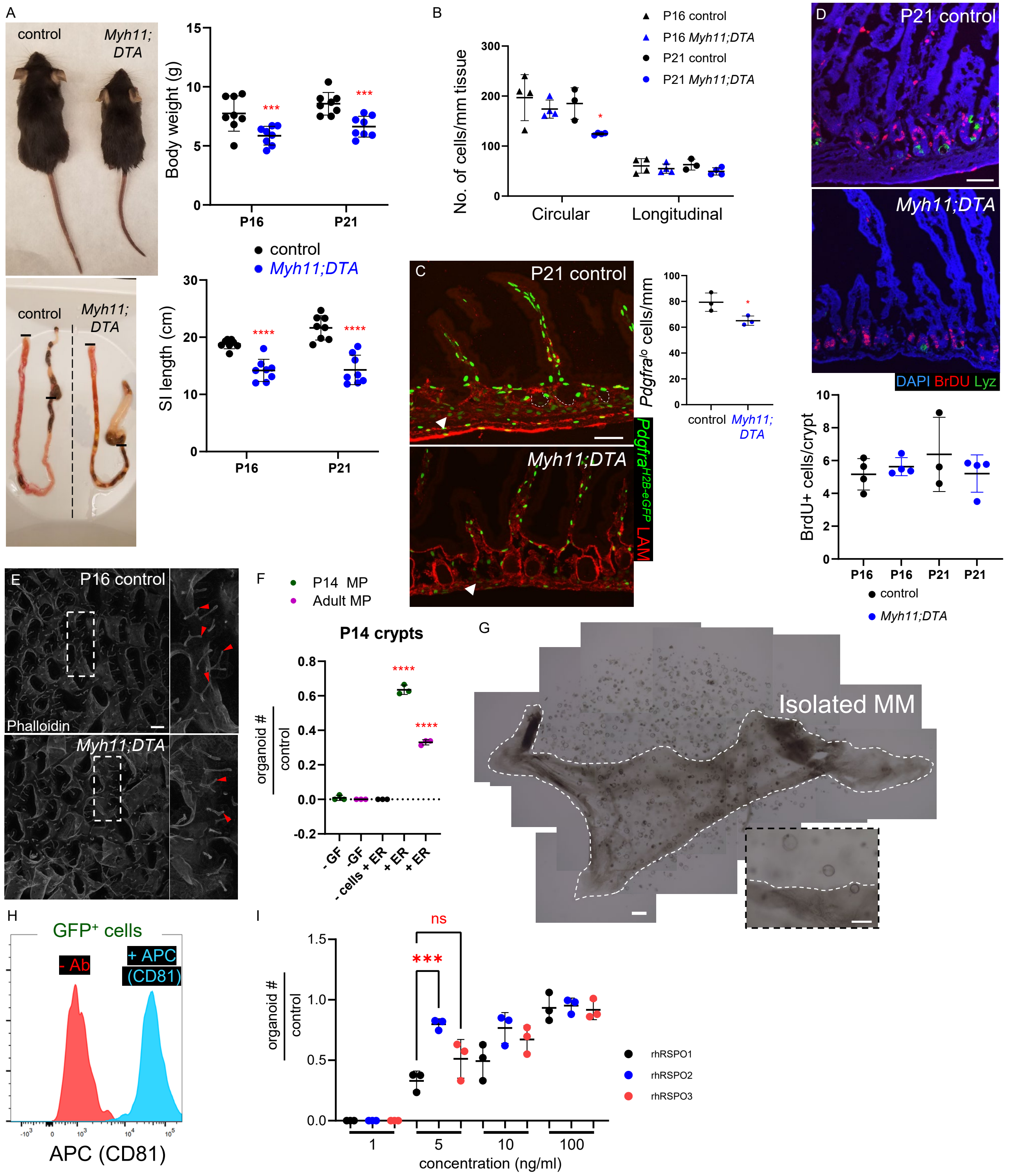
