## Supplementary material for "Delineation and birth of a layered intestinal stem cell niche": Figure S2

A

**Combined datasets:** Unfractionated mesenchyme (P1, P4, P5, P9, P14) and GFP+ cells from *Pdgfra*[H2B-eGFP] intestines (P2, P5, P14)

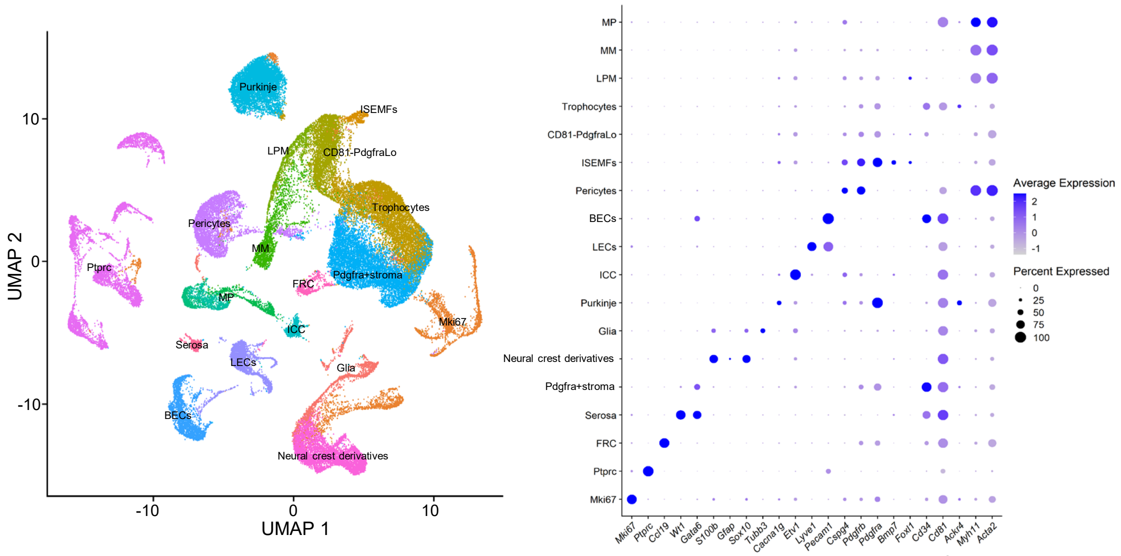


B

CD45/*Ptprc*+ Cells

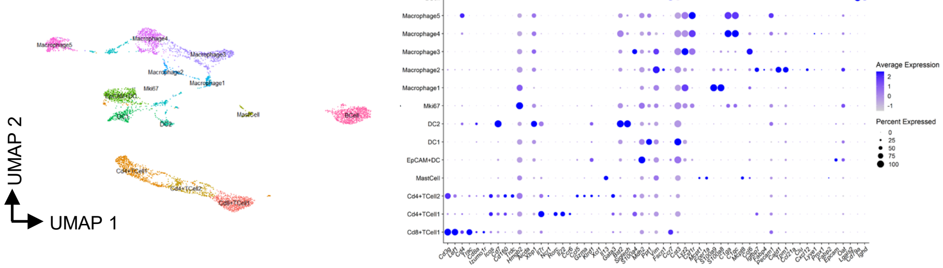


C

% *Mki67*+ cells

| | P1/P2 | P4/P5 | P9 | P14 |
|---|---|---|---|---|
| NCC derivatives: | 45% | 29% | 17% | 7% |
| Pericytes: | 10% | 11% | 5% | 3% |
| Purkinje: | 26% | 12% | 10% | 4% |
| Pdgfra+stroma: | 9% | 8% | 4% | 1% |
| ICC: | 7% | 9% | 3% | 3% |
| Trophocytes: | 16% | 10% | 5% | 2% |
| CD81-PdgfraLo: | 8% | 13% | 15% | 6% |
| Ptprc: | 0% | 4% | 2% | 0% |

D

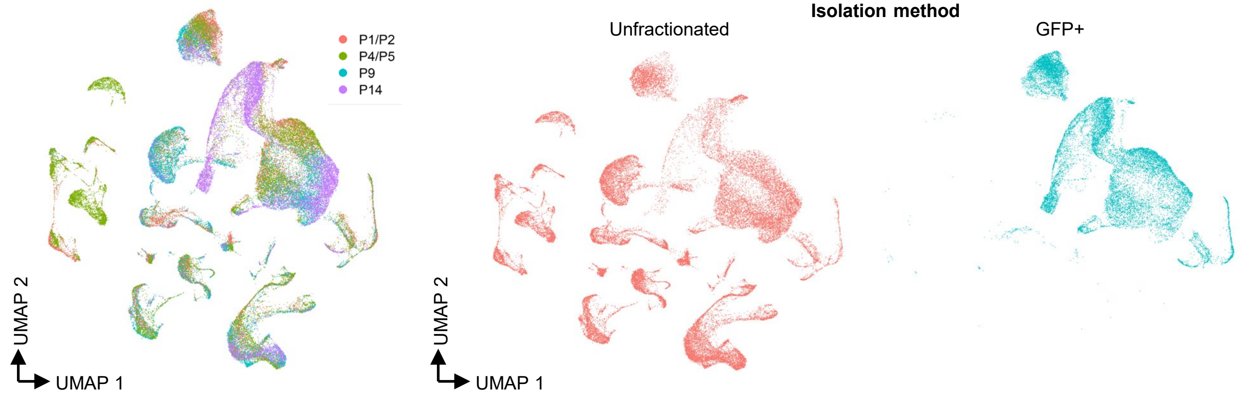


E

| | Glia | ISEMFs | Trophocytes | CD81-PdgfraLo | **LPM** | **MM** | Nog+MP | Pcp4+MP | Osr1+MP | ICC | Purkinje | Pdgfra+stroma | BECs | LECs | Pericytes | Ptprc | NCC Derivatives | FRC | Serosa |
|---|---|---|---|---|---|---|---|---|---|---|---|---|---|---|---|---|---|---|---|
| P1/P2: | 281 | 398 | 2082 | 168 | **17** | **84** | 223 | 435 | 160 | 236 | 2702 | 1447 | 248 | 296 | 412 | 1336 | 370 | 105 | 134 |
| P4/P5: | 1095 | 97 | 2526 | 254 | **90** | **274** | 215 | 147 | 147 | 276 | 1695 | 2102 | 1123 | 1234 | 680 | 4164 | 1520 | 204 | 241 |
| P9: | 605 | 41 | 1307 | 95 | **19** | **89** | 245 | 172 | 211 | 212 | 1250 | 2440 | 814 | 721 | 2652 | 209 | 1703 | 122 | 116 |
| P14: | 239 | 66 | 2092 | 3102 | **935** | **1436** | 138 | 60 | 64 | 61 | 1059 | 2691 | 366 | 271 | 484 | 22 | 1812 | 188 | 93 |
