## Supplementary material for "Delineation and birth of a layered intestinal stem cell niche": Figure S3

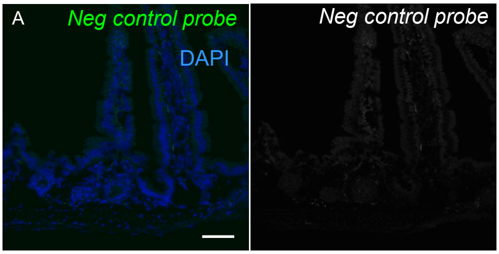


B

| | | *Nog* rep 1 | *Nog* rep 2 | *Pcp4* rep 1 | *Pcp4* rep 2 | *Osr1* rep 1 | *Osr1* rep 2 |
|---|---|---|---|---|---|---|---|
| | MM | 0% (15) | 5% (18) | 9% (11) | 0% (19) | 5% (17) | 10% (19) |
| | Submucosa | 3% (116) | 2% (93) | 6% (179) | 1% (163) | 2% (146) | 12% (126) |
| Circular | Superficial | **41% (107)** | **61% (77)** | 17% (75) | 8% (82) | 3% (94) | 13% (60) |
| Circular | Deep | 2% (276) | 1% (266) | **77% (299)** | **65% (263)** | 4% (245) | 4% (295) |
| | Plexus | 29% (17) | 27% (22) | 11% (17) | 21% (14) | 0% (8) | 15% (13) |
| | Long | 2% (110) | 1% (122) | **47% (151)** | **38% (145)** | **87% (115)** | **70% (136)** |

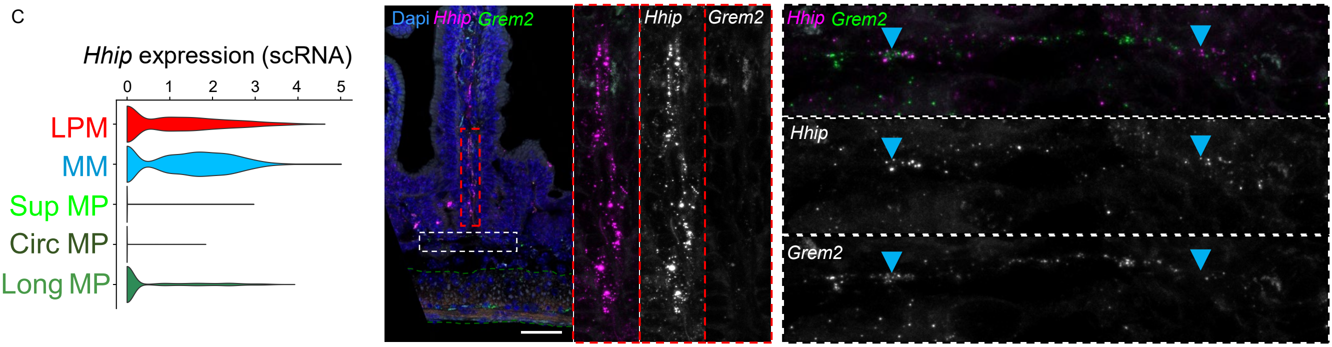


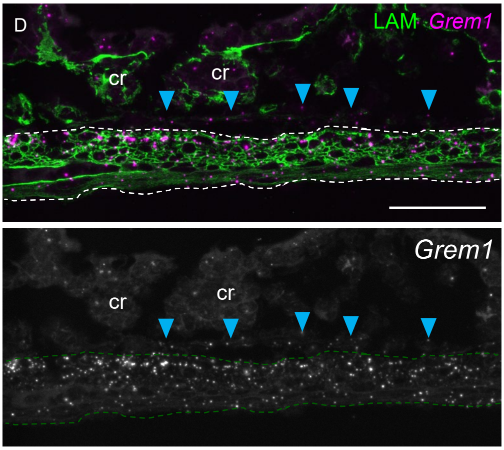


| | | *Grem1* rep 1 | *Grem1* rep 2 | *Grem2* rep 1 | *Grem2* rep 2 |
|---|---|---|---|---|---|
| | MM | **80% (15)** | **83% (18)** | **91% (22)** | **81% (22)** |
| | Submucosa | 41% (101) | 26% (182) | 12% (196) | 4% (162) |
| Circular | Superficial | **97% (78)** | **90% (142)** | 24% (103) | 38% (100) |
| Circular | Deep | **83% (243)** | **76% (129)** | 10% (277) | 8% (331) |
| | Plexus | 0% (6) | 11% (27) | **77% (22)** | **48% (31)** |
| | Long | **61% (170)** | **73% (152)** | 15% (118) | 11% (20) |

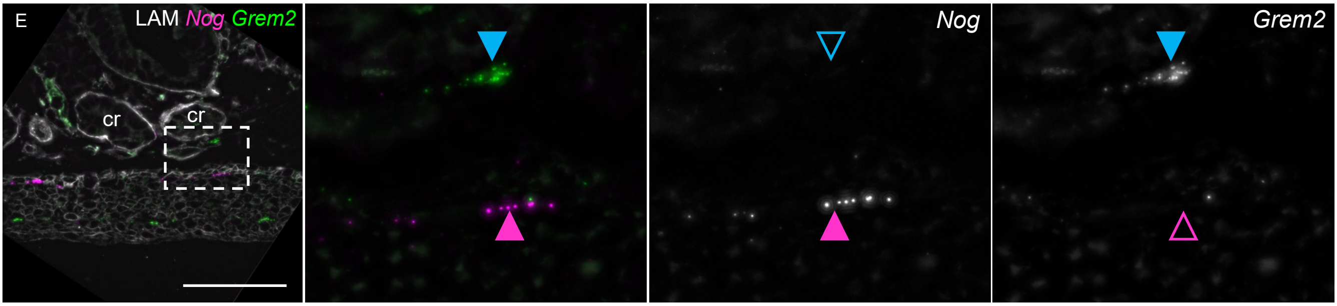
